# Association with TFIIIC limits MYCN localization in hubs of active promoters and chromatin accumulation of non-phosphorylated RNA Polymerase II

**DOI:** 10.1101/2023.11.18.567687

**Authors:** Raphael Vidal, Eoin Leen, Steffi Herold, Mareike Müller, Daniel Fleischhauer, Christina Schülein-Völk, Dimitrios Papadopoulos, Isabelle Röschert, Leonie Uhl, Carsten P. Ade, Peter Gallant, Richard Bayliss, Martin Eilers, Gabriele Büchel

## Abstract

MYC family oncoproteins regulate the expression of a large number of genes and broadly stimulate elongation by RNA polymerase II. While the factors that control the chromatin association of MYC proteins are well understood, much less is known about how interacting proteins mediate MYC’s effects on transcription. Here we show that TFIIIC, an architectural protein complex that controls the three-dimensional chromatin organization at its target sites, binds directly to the amino-terminal transcriptional regulatory domain of MYCN. Surprisingly, TFIIIC has no discernible role in MYCN-dependent gene expression and transcription elongation. Instead, MYCN and TFIIIC preferentially bind to promoters with paused RNAPII and globally limit the accumulation of non-phosphorylated RNAPII at promoters. Consistent with its ubiquitous role in transcription, MYCN broadly participates in hubs of active promoters. Depletion of TFIIIC further increases MYCN localization to these hubs. This increase correlates with a failure of the nuclear exosome and BRCA1, both of which are involved in nascent RNA degradation, to localize to active promoters. Our data suggest that MYCN and TFIIIC exert an censoring function in early transcription that limits promoter accumulation of inactive RNAPII and facilitates promoter-proximal degradation of nascent RNA.

## Introduction

The MYC family of proto-oncogenes is at the epicenter of cellular regulatory networks that govern cell growth, proliferation, and differentiation (Dang, 2012, Kress, Sabo et al., 2015). The three MYC paralogs (MYC, MYCN, and MYCL) are central players in normal development and tissue homeostasis, and when dysregulated, fuel many of the processes that are hallmarks of cancer (Dhanasekaran, Deutzmann et al., 2022, Hanahan, 2022). Among them, MYCN has attracted attention for its causal role in the development of neuroblastoma and other childhood tumors (Rickman, Schulte et al., 2018).

Both MYC and MYCN bind to virtually all active promotors and profoundly alter the dynamics of RNA polymerase II (RNAPII) transcription, with an increase in pause release and elongation being most apparent (Herold, Kalb et al., 2019, Walz, Lorenzin et al., 2014). One consequence of these changes are alterations in expression of a broad range of target genes (Dhanasekaran, Hansen et al., 2023). Unrelated to those changes in gene expression, MYC and MYCN also control RNAPII function to limit the accumulation of R-loops, to facilitate promoter-proximal double-strand break repair and to co-ordinate transcription elongation with DNA replication (Papadopoulos, Uhl et al., 2023). Which of these effects are critical for the oncogenic functions of MYC is an open question and consequently the partner proteins via which MYC proteins alter RNAPII dynamics are under intense investigation (Baluapuri, Hofstetter et al., 2019, Baluapuri, Wolf et al., 2020, Buchel, Carstensen et al., 2017, Das, Lewis et al., 2022, Heidelberger, Voigt et al., 2018, Kalkat, Resetca et al., 2018, Lourenco, Resetca et al., 2021, Oksuz, Henninger et al., 2023). Direct interactions of MYC and MYCN with MAX, WDR5 and MIZ1 control the localization of MYC on chromatin (Blackwood & Eisenman, 1991, Blackwood, Lüscher et al., 1992, Thomas, Wang et al., 2015, Vo, Wolf et al., 2016, Walz et al., 2014). MYC-dependent effects on RNAPII elongation involves the transfer of elongation factors SPT5 and PAF1c from MYC onto RNAPII (Baluapuri et al., 2019, Endres, Solvie et al., 2021, Jaenicke, von Eyss et al., 2016). Both MYC and MYCN also interact with and activate topoisomerases I and II, suggesting that MYC/N-dependent pause release also involves the relieve of torsional stress that builds up during early transcription (Das, Kuzin et al., 2022).

Intriguingly, both MYC and MYCN form prominent complexes with TFIIIC and mapping experiments for MYCN show that the amino terminal transcription-regulatory domain is required for the interaction (Buchel et al., 2017, Heidelberger et al., 2018). TFIIIC was first identified as a general transcription factor for RNA polymerase III (RNAPIII) (Orioli, Pascali et al., 2012). Surprisingly, TFIIIC also binds to thousands of genomic sites that are not shared with RNAPIII and are called <u>e</u>xtra <u>T</u>FIII<u>C</u> (ETC) sites (Noma, Cam et al., 2006). Many ETC sites localize to RNAPII-transcribed promoters and such sites are often juxtaposed to MYCN binding sites (Buchel et al., 2017, Moqtaderi, Wang et al., 2010, Noma et al., 2006, Oler, Alla et al., 2010). TFIIIC is an architectural protein that can affect the three-dimensional chromatin organization at its binding sites (Noma et al., 2006, Van Bortle & Corces, 2013). Functionally TFIIIC can act as insulator that blocks the spread of chromatin states (Raab, Chiu et al., 2012), affect cohesin loading (Buchel et al., 2017), and bind at the borders of topologically <u>a</u>ssociating <u>d</u>omains (TADs) (Van Bortle, Nichols et al., 2014).

Intriguingly, TFIIIC compartmentalizes RNAPII promoters and gene expression. Specifically, the effects of TFIIIC on the expression of E2F-dependent and neuronal genes correlate with its effects on the three-dimensional chromatin architecture. In the case of neuronal genes, TFIIIC prevents the localization of activity-dependent genes to sites of active transcription before stimulation (Crepaldi, Policarpi et al., 2013, Ferrari, de Llobet Cucalon et al., 2020, Policarpi, Crepaldi et al., 2017). Similarly, TFIIIC together with the <u>a</u>ctivity-<u>d</u>ependent <u>n</u>euro<u>p</u>rotector homeobox protein (ADNP) controls the three-dimensional architecture of cell cycle genes (Ferrari et al., 2020). Our previous experiments had suggested that TFIIIC may have a role in MYCN-dependent control of RNAPII function and we here show that MYCN and TFIIIC together have an unexpected censoring function that excludes non-functional RNAPII from hubs of active promoters.

## Results

### TFIIIC interacts directly with the amino-terminus of MYCN

TFIIIC is present in MYCN immunoprecipitates and the amino terminal region of recombinant MYCN binds to TFIIIC in pull-down experiments performed with cell lysates (Buchel et al., 2017). To test whether MYCN and TFIIIC interact directly, we expressed the six subunits of TFIIIC together with a FLAG-tagged MYCN construct comprising amino acids 2-137 using a recombinant baculovirus to infect insect cells. Performing pulldown experiments with cell lysates and immobilized FLAG-tagged MYCN, we recovered all six subunits of the TFIIIC complex in the eluate along with the FLAG-tagged MYCN construct, demonstrating that TFIIIC can bind to the amino terminal region of MYCN (Figure 1A). TFIIIC is composed of two subcomplexes, designated TauA (τA) and TauB (τB), each comprising three of the six subunits. We first attempted co-expression of FLAG-MYCN with individual subcomplexes, but the τB complex was too unstable to be isolated, and so we focused on the stable τA complex. To test whether MYCN binds directly to τA, we purified MYCN and the τA complex separately (Figure S1A, B) and then mixed the purified preparations (Figure 1B-D). In gel filtration experiments, the isolated MYCN protein eluted with a molecular weight of around 50 kDa, whereas a fraction of MYCN eluted with a much larger molecular weight together with the τA complex when both were mixed (Figure 1B-D). Native-mass spectrometry (Tamara, den Boer et al., 2022) showed the molecular weight of this complex to be 204.7 kDa, very close to the sum of the molecular weights of a 1:1:1:1 complex (Figure S1C, D). We concluded that the MYCN amino terminal region and the τA subcomplex of TFIIIC form a stable complex with each other in solution.

**Figure 1.**
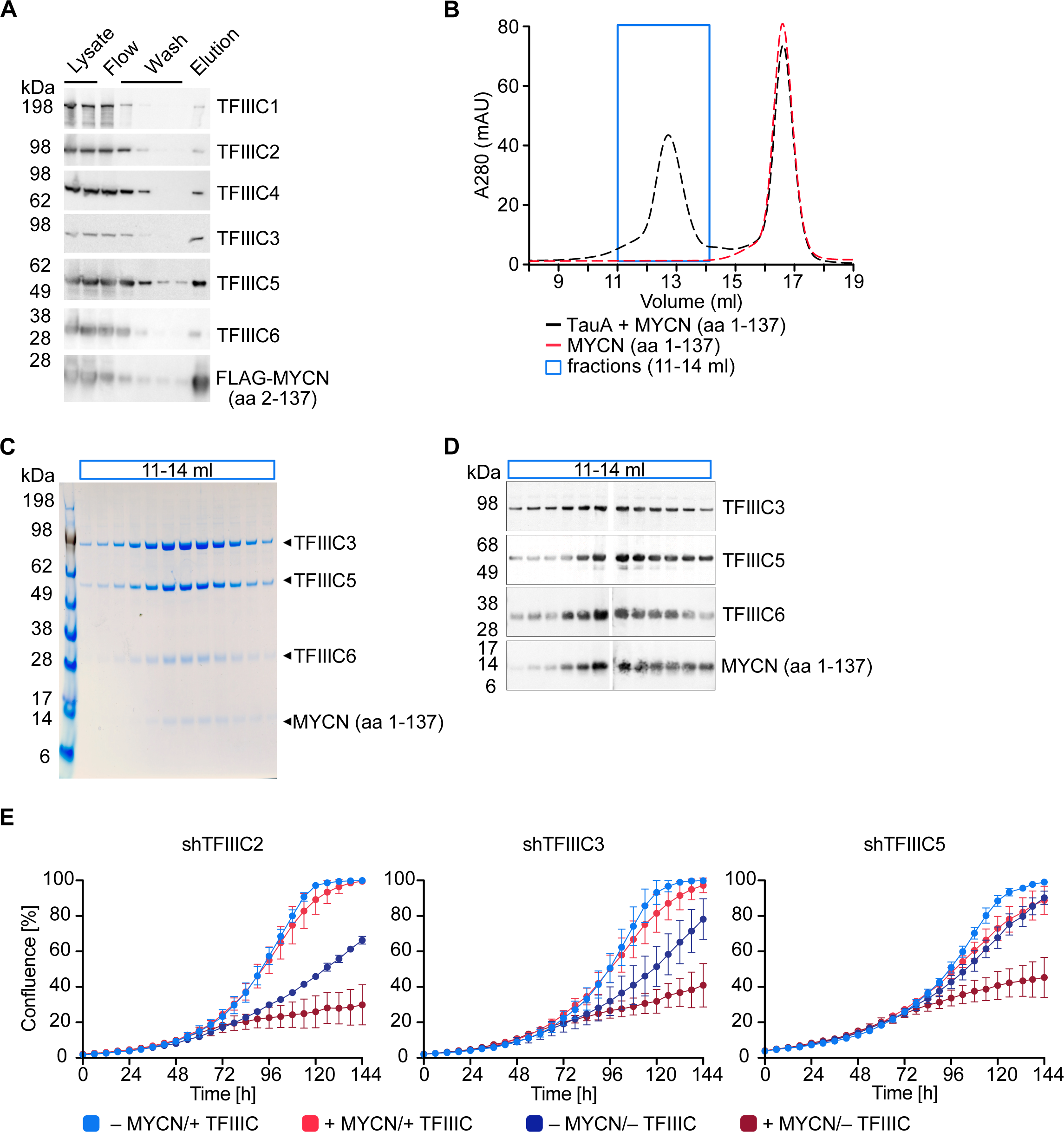
TFIIIC directly interacts with MYCN. **A.** Immunoblots showing levels of FLAG-tagged MYCN (amino acids 2-137) and the six subunits of the TFIIIC complex after a pulldown assay using anti-FLAG affinity columns. Multiple columns labelled “Wash” represent the sequential washings (n = 2). **B.** Size exclusion chromatography graph of MYCN (aa 1-137)/1A (black trace) or MYCN alone (red trace). The blue box marks the fractions used for panel C and D (n = 2). **C.** Coomassie staining of fractions of the MYCN (aa 1-137)/1A complex (fractions marked with blue box in panel B). **D.** Immunoblot of fractions of the MYCN (aa 1-137)/1A complex (fractions marked with blue box in panel B). **E.** Growth curve (measured as % confluence) of SH-EP-MYCN-ER cells expressing Dox-inducible shRNA targeting *TFIIIC2*, *TFIIIC3* or *TFIIIC5* under the indicated conditions. Data show mean ± standard deviation (SD) (n = 3).

As described in the introduction, TFIIIC is both a general transcription factor for RNAPIII and has been implicated in regulation of RNAPII-dependent transcription. To test whether TFIIIC is required for the proliferation of neuroblastoma cells, we individually depleted three of its subunits, TFIIIC2, TFIIIC3 and TFIIIC5, by stable expression of doxycycline-inducible shRNAs. Controls confirmed that each shRNA efficiently depleted its target protein (Figure S1E). To test both for a general requirement of each subunit in neuroblastoma cell proliferation and for MYCN-specific effects, we expressed each shRNA in SH-EP-MYCN-ER cell. SH-EP cells express endogenous MYC and are engineered to stably express a MYCN-ER chimera (Herold et al., 2019). Activation of the MYCN-ER by addition of 4-hydroxytamoxifen (4-OHT) suppresses the expression of endogenous MYC (Figure S1F), in effect causing a switch from MYC to MYCN expression. Under control conditions, depletion of TFIIIC2 or TFIIIC3 attenuated proliferation, whereas depletion of the small TFIIIC5 subunit had little effect. Addition of 4-OHT had little effect by itself, but greatly enhanced the inhibitory effect of depletion of each subunit (Figure 1E), suggesting that TFIIIC has both general and MYCN-specific roles in neuroblastoma cell proliferation.

### TFIIIC limits promoter binding of non-phosphorylated RNAPII in MYCN-expressing cells

The proximity of TFIIIC binding sites to MYCN binding sites at many promoters transcribed by RNAPII prompted us to explore the role of TFIIIC in MYCN-driven transcription (Buchel et al., 2017). Using an antibody that detects total RNAPII, we had previously shown that activation of MYCN in SH-EP-MYCN-ER cells impacts RNAPII in two ways: First, MYCN promotes pause release and MYCŃs effect on elongation correlates with its effects on gene expression (Herold et al., 2019, Rahl, Lin et al., 2010, Walz et al., 2014). At the same time, activation of MYCN uniformly reduces RNAPII occupancy at all active promoters; this correlates with a reduced accumulation of promoter-proximal R-loops, suggesting that MYCN limits the accumulation of RNAPII at promoters that is impaired in productive elongation and/or splicing (Herold et al., 2019).

Previous experiments using antibodies directed against total RNAPII had yielded results that varied among different antibodies ( (Buchel et al., 2017) and RV, unpublished). To directly compare the effects of MYCN and TFIIIC on promoter-bound RNAPII with those on elongating RNAPII in a side-by-side manner, we conducted parallel ChIP-sequencing (ChIP-seq) analyses using antibodies that specifically recognize non-phosphorylated and Ser2-phosphorylated (pSer2) RNAPII, respectively. Visual inspection of multiple individual genes (Figure 2A) and global analyses (Figure 2B,C and Figure S2A,B) showed that activation of MYCN caused a global decrease in promoter association of non-phosphorylated RNAPII. A control ChIP-seq experiment established that addition of 4-OHT had no such effect in SH-EP cells that do not express a MYCN-ER chimera, demonstrating that the reduction is due to activation of MYCN (Figure S2C). At the same time, MYCN promoted an increase in transcription elongation as best documented by pSer2-RNAPII occupancy at the transcription end site (TES) (Figures 2A, C). This increase was most pronounced on MYCN-activated genes. Comparison with RNA-sequencing (RNA-seq) data showed that it correlated closely with MYCN-dependent changes in gene expression (Figure 2D). Depletion of TFIIIC5 abrogated the MYCN-dependent decrease in chromatin association of non-phosphorylated RNAPII but had no effect on MYCN-dependent changes in transcription elongation (Figure 2A-C). In many experimental systems, MYC and MYCN effects on elongation parallel closely with the corresponding changes in gene expression. Consistent with these observations, RNA-seq showed that neither depletion of TFIIIC3 nor of TFIIIC5 had significant effects on either basal gene expression or MYCN-dependent changes in steady-state mRNA levels (Figures 2E and Figure S2D). Finally, we aimed to understand whether the localization of TFIIIC and MYCN at promoters is linked to the dynamics of RNAPII at promoters and performed ChIP-seq of TFIIIC5 in SH-EP-MYCN-ER cells in the presence of 5,6-Dichlorobenzimidazole-1-β-D-ribofuranoside (DRB), which stabilizes paused RNAPII and prevents pause-release of RNAPII (Mancebo, Lee et al., 1997). Inspection of individual genes (Figure S2E) and average density plots of all active promoters (Figure 2F) showed that activation of MYCN globally enhanced TFIIIC5 association with regions surrounding active transcription start sites, consistent with previous observations (Buchel et al., 2017). Addition of DRB increased TFIIIC5 association with active promoters and the combination of MYCN activation and DRB had a strong additive effect, arguing that MYCN preferentially recruits TFIIIC5 to promoters with paused RNAPII.

**Figure 2.**
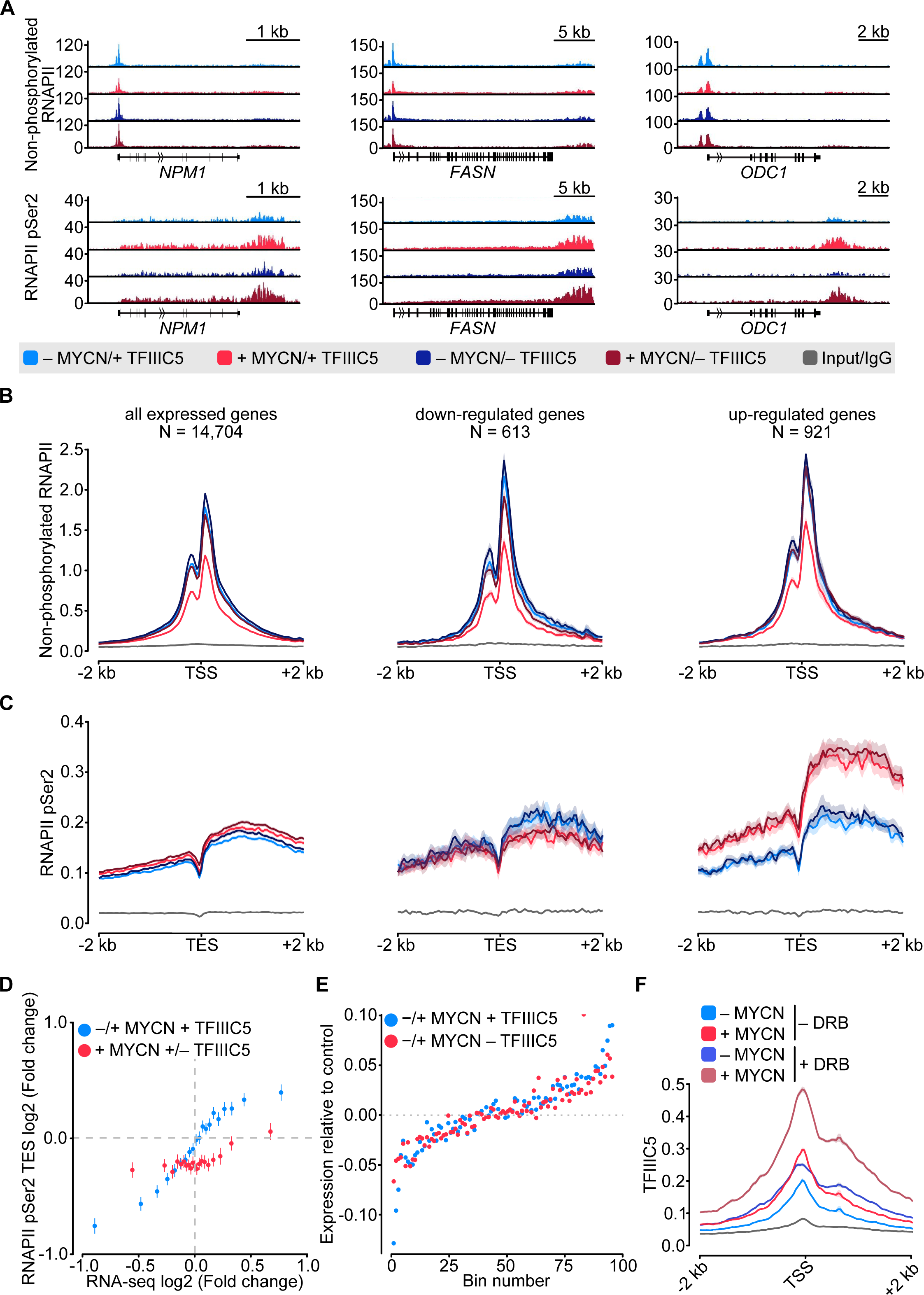
MYCN and TFIIIC antagonize accumulation of non-phosphorylated RNAPII. **A.** Browser tracks for non-phosphorylated RNAPII (top) and RNAPII pSer2 (bottom) ChIP-Rx at the indicated gene loci. SH-EP-MYCN-ER cells were treated with Dox (1 µg/ml, 48 h) and/or 4-OHT, respectively. EtOH was used as control. **B.** Average density plot of ChIP-Rx signal for non-phosphorylated RNAPII. Data show mean (line) ± standard error of the mean (SEM indicated by the shade) of different gene sets based on an RNA-seq of SH-EP-MYCN-ER cells ± 4-OHT. The y-axis shows the number of spike-in normalized reads and it is centered to the TSS ± 2 kb. N = number of genes in the gene set defined in the methods (n = 2). **C.** Density plot of ChIP-Rx signal for RNAPII pSer2 as described for panel B. The signal is centered to the TES ± 2 kb (n = 2). **D.** Average bin dot plot showing fold change for RNAPII pSer2 ChIP-Rx reads over TES ± 2 kb and RNA-seq of SH-EP-MYCN-ER for the same genes ± MYCN + TFIIIC5 (blue) or + MYCN ± TFIIIC5 (red). The plot shows 20 bins representing a total of 13,239 and 12,330 genes for ± MYCN + TFIIIC5 and + MYCN ± TFIIIC5 datasets, respectively (n = 3 for RNA-seq, n = 2 pSer2 RNAPII ChIP-Rx). **E.** Average bin dot plot for RNA-seq of SH-EP-MYCN-ER showing log2 mRNA expression normalized by control per bin. Cells were treated with 1 µg/ml Dox (“– TFIIIC5”, 48 h) and/or 4-OHT (“+ MYCN”, 4 h) or EtOH as control. Expression was normalized by its control. Each bin represents 150 genes of a total of 14,085 genes. Dotted line marks the relative expression at 0 (n = 3). **F.** Density plot of ChIP-Rx signal for TFIIIC5. Data show mean (line) ± SEM (shade) for 14,722 genes. The signal is centered to the TSS ± 2 kb (n = 2).

### MYCN takes part in three-dimensional networks of active promoters

The effects of TFIIIC on the expression of E2F-dependent and neuronal genes correlate with its effects on the three-dimensional chromatin architecture (see Introduction). This raised the possibility that complex formation of TFIIIC similarly affects three-dimensional chromatin interactions of MYCN. To test this hypothesis, we used <u>p</u>hosphorylated linker HiChIP (pLHiChIP), a modification of the HiChIP protocol, which identifies pairs of DNA loci that are brought into close spatial proximity to a specific protein (Figure S3A) (Mumbach, Rubin et al., 2016). We initially performed pLHiChIP for MYCN in SH-EP-MYCN-ER neuroblastoma cells (Figure 3A). ChIP experiments measuring the occupancy of multiple promoters bound by both MYCN and MYC showed that the chromatin association of MYCN is much greater than that of MYC, allowing analysis of MYCN function on chromatin with little interference from MYC (Figure S3B). Visual inspection of the MYCN pLHiChIP showed that thousands of pairs of MYCN binding sites are in close spatial proximity with each other (Figure 3A). Appropriate quality controls established the validity of these results: For example, the Hi-C module of pLHiChIP protocol yielded a high percentage of valid interactions pairs (Figure S3C, D). Relative to the Hi-C input, the MYCN HiChIP is strongly enriched for interactions that connected two MYCN-binding sites with each other, confirming the specificity of the signal (Figure S3E).

**Figure 3.**
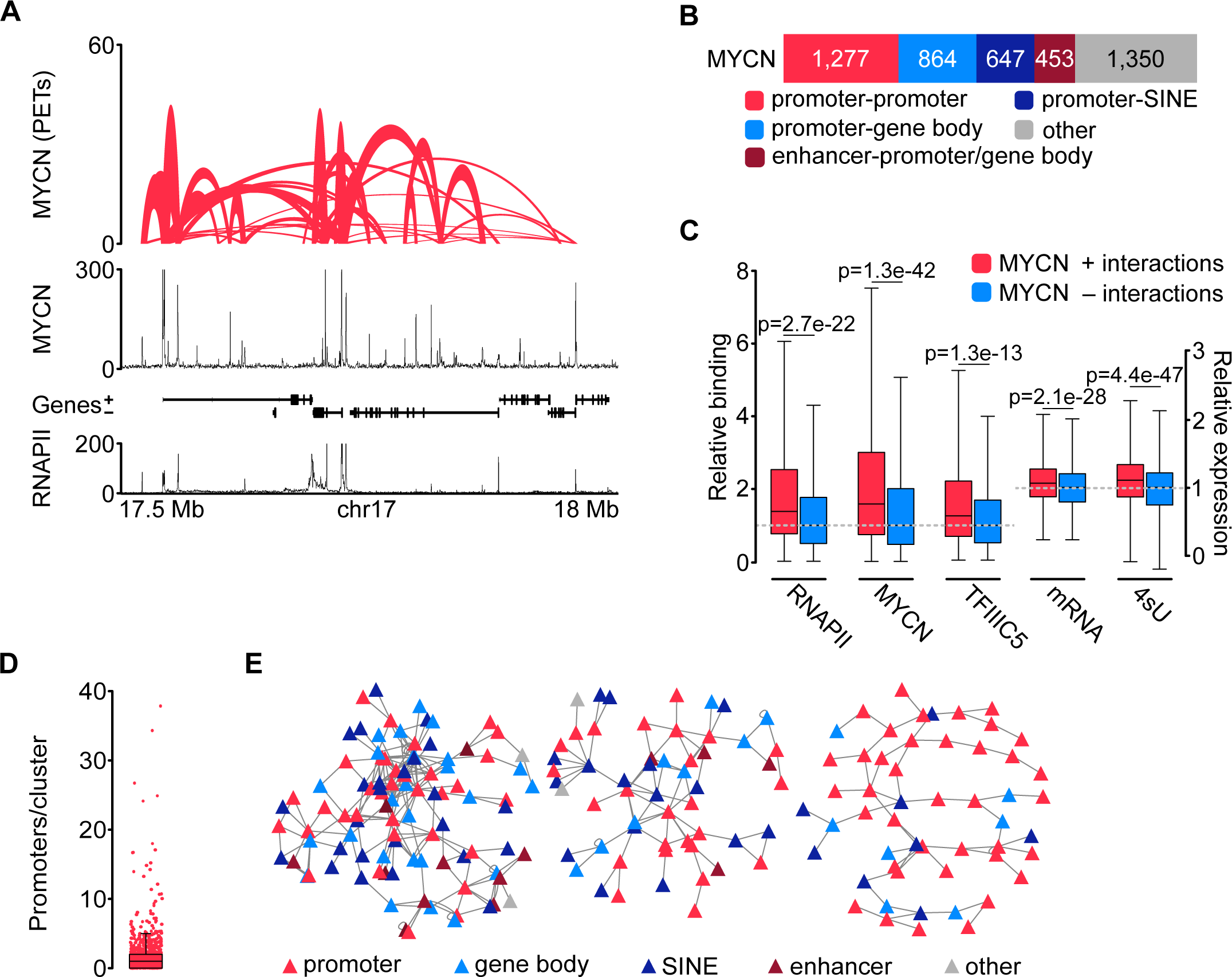
MYCN is part of three-dimensional promoter hubs. **A.** Top: Representative browser track of MYCN three-dimensional chromatin interactions. Height shows the number of <u>p</u>aired <u>e</u>nd tags (PETs) indicating the interaction intensity and the width of the line shows the start and end positions of each anchor. Middle and bottom: Browser tracks showing the number of reads of MYCN and total RNAPII ChIP-Rx, respectively. Unless stated, all experiments were performed in SH-EP-MYCN-ER cells treated with 4-OHT (200 nM, 4 h). The ruler at the bottom shows the genomic coordinates. (n = 3 independent biological replicates for MYCN pLHiChIP; n = 2 for RNAPII ChIP-Rx). **B.** Bar chart listing functional annotations of all binary MYCN interactions (N = 4,591; N indicates total number). **C.** Boxplots showing relative binding of the indicated proteins (RNAPII, MYCN, TFIIIC5) to promoter regions or expression levels of the corresponding genes (mRNA by RNA-seq; 4sU by 4sU-seq). Red boxes: Genes bound by MYCN and part of MYCN-hubs; Blue boxes: Genes bound by MYCN that are not part of MYCN-hubs. Each pair was normalized to the median of the corresponding “blue” gene set. p-values were obtained by pairwise comparisons using Student’s *t-*test. (n = 2 for TFIIIC5 and RNAPII ChIP-Rx). **D.** Boxplot showing the number of promoters in each cluster, with each red dot representing one cluster. **E.** Network reconstruction of the three biggest clusters based on MYCN pLHiChIP interactions. Each anchor is represented by a node (“triangle”) and the lines show interactions between the anchors. The colors are indicating the different functional annotation.

The pLHiChIP data analysis revealed a total of 4,591 distinct binary interactions involving MYCN (Figure 3B). To facilitate a comprehensive functional analysis of MYCN anchor sites, we performed ChIP-seq for RNAPII from SH-EP-MYCN-ER cells and integrated the HiChIP data with these data (Figure 3A), along with other relevant annotations, such as SINE elements. Remarkably, our analysis revealed that 1,277 out of the 4,591 MYCN anchors (28%) were positioned on RNAPII promoters (Figure 3B). Furthermore, 864 out of the 4,591 MYCN interactions (19%) exhibited connections between promoters and either exonic or intronic sequences, consistent with previous observations showing that MYCN binding sites are often located within transcribed regions (Buchel et al., 2017). Additionally, MYCN interactions were observed between promoters and SINE repetitive elements (647/4,591; 14%) including interactions with tRNAs (Figure S3F). These connections are likely shared with TFIIIC (see below). We also observed interactions between promoters or gene bodies and enhancers, accounting for 453 out of the 4,591 interactions (10%).

We used the RNAPII and published MYCN (Herold et al., 2019) ChIP-seq data as well as sequencing datasets of mRNA (Buchel et al., 2017) and nascent (4sU-labeled) (Papadopoulos, Solvie et al., 2022) RNA to identify the specific properties of promoters within three-dimensional MYCN interactions. This showed that significantly more MYCN and RNAPII bound to promoters that participated in such interactions and that these promoters were more active than MYCN-bound promoters without such interactions, although the latter difference was small (Figure 3C). We next used the binary interactions as a starting point to reconstruct interaction networks. This showed that MYCN participated in three-dimensional promoter hubs of different sizes (Figure 3D), with the largest connecting up to 34 promoters and 10 enhancers (Figure 3E). This number of promoters is consistent with published estimates (Palacio & Taatjes, 2022). Functional annotation of the promoters that are contained in these large hubs showed that they are highly enriched in genes encoding ribosomal proteins and ribosome biogenesis genes transcribed by RNAPII (Figure S3G). We concluded from the data that MYCN participates in three-dimensional hubs of highly active promoters; we will use the term “promoter hubs” (Lim & Levine, 2021) for these structures.

### Binding to TFIIIC antagonizes MYCN localization in promoter hubs

To determine whether TFIIIC participates in three-dimensional chromatin structures, we performed pLHiChIP using a previously validated antibody against TFIIIC5 (Buchel et al., 2017). In these experiments, thousands of pairs of TFIIIC binding sites yielded a robust signal, demonstrating that they are in close spatial proximity with each other (Figure 4A). Specifically, these analyses identified a total of 3,499 binary non-contiguous interactions for TFIIIC. Comparison to RNAPII ChIP-seq data showed that 31% of these interactions connected a promoter or the gene body of an RNAPII-transcribed gene to a SINE repetitive element (Figure 4B). Importantly, 1,107 out of 4,591 MYCN interactions shared one (1,075) or both (32) anchors with TFIIIC-containing interactions arguing that both proteins participate in common hubs (Figure 4C). This is consistent with our previous data, which showed that many promoters contain a TFIIIC-bound site close to a MYCN-bound E-box element (Buchel et al., 2017). A significant percentage of the interactions that are shared between MYCN and TFIIIC contained an E-box in one anchor and A- or B-boxes, which are bound by TFIIIC, in the other anchor sequence, arguing that MYCN and TFIIIC can participate in the same interactions (Figure S4A). Most interactions that are shared by MYCN and TFIIIC connected promoters to other promoters, to gene bodies or to SINE elements. Conversely, TFIIIC5 anchors without overlapping MYCN loops (“TFIIIC5 only”) did not show an enrichment for promoter interactions (Figure 4D). We concluded that both MYCN and TFIIIC5 are present in promoter hubs. To determine how TFIIIC affects MYCN involvement in promoter hubs, we used SH-EP-MYCN-ER neuroblastoma cells that express a Dox-inducible shRNA targeting *TFIIIC5* (Figure S1E) and performed <u>s</u>pike-in <u>p</u>hosphorylated linker HiChIP (spLHiChIP), a spike-in variation of pLHiChIP that allows quantitative comparisons between different samples. These experiments showed that depletion of TFIIIC5 strongly increased the number of chromatin interactions of MYCN, arguing that association with TFIIIC antagonizes MYCN participation in promoter hubs (Figure 4E, F and Figure S4B). HiChIP experiments performed for TFIIIC5 showed that expression of MYCN moderately and blockade of pause release by DRB strongly decreased the frequency of three-dimensional interactions of TFIIIC5 (Figures 4G, H). Stratification showed that interactions of TFIIIC5 with promoters, gene bodies and SINE elements were all decreased upon incubation of cells with DRB (Figure S4C), arguing that MYCN/TFIIIC5-bound genes are not part of three-dimensional promoter hubs.

**Figure 4.**
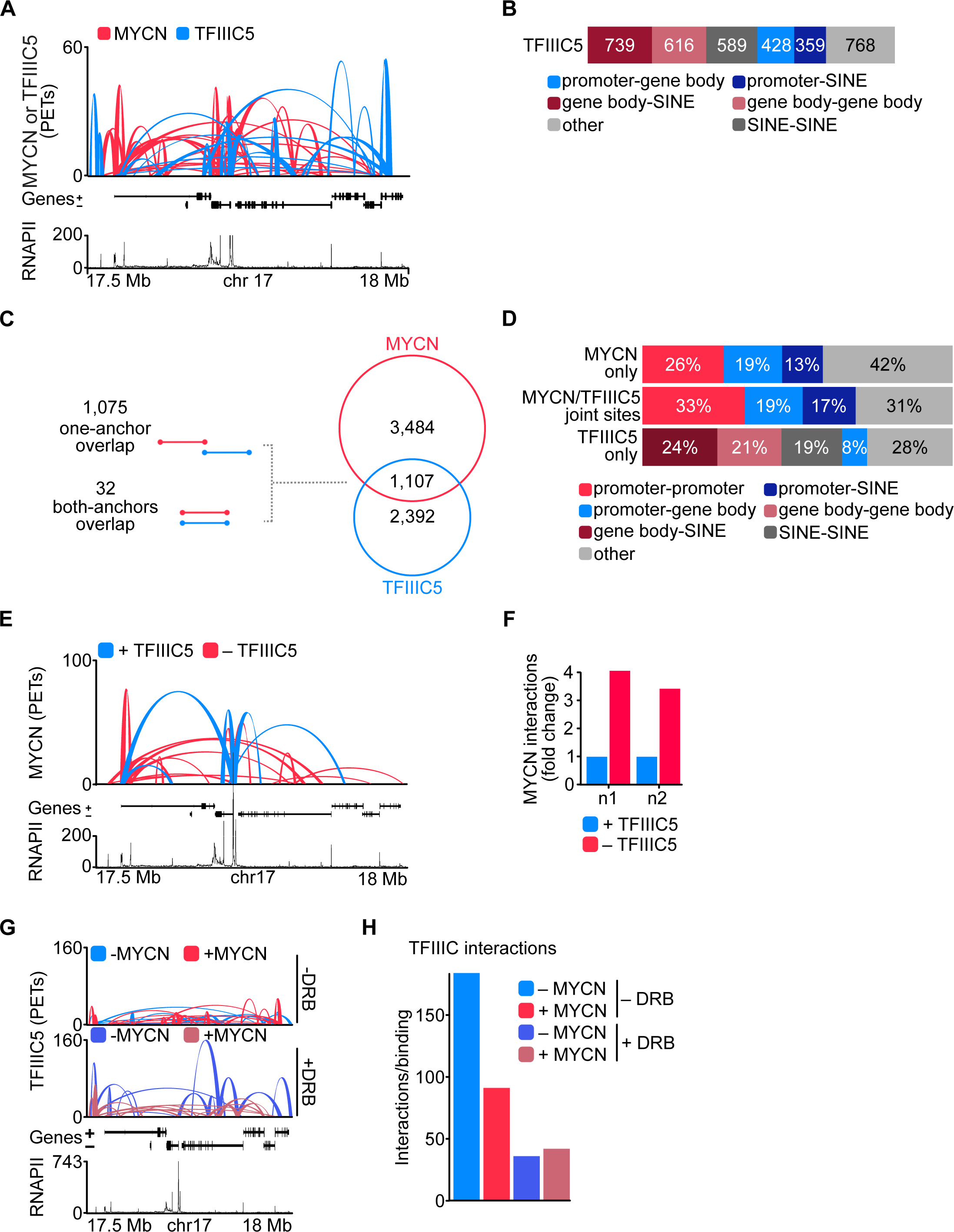
TFIIIC antagonizes MYCN participation in promoter hubs. **A.** Representative example of pLHiChIP track for MYCN (red) and TFIIIC5 (blue) interactions (conventions as in Figure 3A) (n = 2). **B.** Bar chart listing the total number of functional annotations for all TFIIIC5 binary interactions (N = 3,499). **C.** Venn diagram showing the number of interactions shared between MYCN and TFIIIC5. The diagram at the left shows the types of overlaps between connections. **D.** Bar chart listing the interactions functional annotations for MYCN anchors not overlapping with TFIIIC5 anchors (“MYCN only”) as well as TFIIIC5 anchors without overlapping MYCN anchors (“TFIIIC5 only”) and their joint anchors. **E.** Representative example of MYCN spLHiChIP track for MYCN interactions in the presence (blue) or absence (red) of TFIIIC5 (n = 2). **F.** Bar graph showing the fold change of all MYCN spLHiChIP interactions comparing “+ TFIIIC5” and “– TFIIIC5” in SH-EP-MYCN-ER cells expressing a Dox-inducible shRNA targeting *TFIIIC5*. n1,2 indicates two independent biological replicates. **G.** Representative example of TFIIIC5 spLHiChIP track without (blue) or with (red) induction of MYCN for SH-EP-MYCN-ER cells (conventions as in Figure 3A). **H.** Bar graph showing the number of TFIIIC5 interactions normalized by the relative binding of TFIIIC5 ChIP-Rx signals for the same coordinates. Coordinates defined as TSS ± 2 kb of 14,722 genes.

### TFIIIC is required for promoter association of factors involved in nascent RNA degradation

Depletion of TFIIIC led to the accumulation of non-phosphorylated RNAPII and an enhanced presence of MYCN in promoter hubs, suggesting that lack of TFIIIC might lead to enhanced or uncontrolled functionality of RNAPII. To ascertain whether TFIIIC is required for functionality of elongating RNAPII, we first performed an rMATS analysis, which analyzes changes in splicing (Shen, Park et al., 2014b, Wang, Pan et al., 2017). This showed that depletion of TFIIIC3 caused significant increases in intron retention and exon skipping, evidence of aberrant splicing (Figure S5A, B). However, most of these alterations occurred in downstream exons, and were not observed for TFIIIC5. Furthermore, expression of MYCN weakened this effect for TFIIIC3 (Figure S5A, B), arguing that it does account for the requirement for TFIIIC function in MYCN-expressing cells.

To explore promoter-proximal events, we made use of the fact that many of the factors that determine premature termination and degradation of nascent RNA have recently been elucidated and initially used PLA assays to survey a series of factors involved in premature termination (Rodriguez-Molina, West et al., 2023). Appropriate controls using immunofluorescence and PLA assays with single antibodies established the specificity of each assay (Figure S5C). Consistent with the ChIP-seq data, activation of MYCN enhanced the proximity between TFIIIC5 and total RNAPII and depletion of TFIIIC5 reduced the signal, confirming the specificity of the assay (Figure S5D). Activation of MYCN and depletion of TFIIIC5 had only weak effects on the proximity of RNAPII with NELFE, which associates with pausing RNAPII. Similarly, depletion of TFIIIC5 either alone or in combination with MYCN activation enhanced the proximity of RNAPII with PP2A, which is recruited to promoters via its interaction with the integrator termination complex (Cossa, Parua et al., 2021, Vervoort, Welsh et al., 2021), and with PNUTS, a targeting subunit of PP1 that is globally involved in termination (Figure 5A) (Cortazar, Sheridan et al., 2019, Estell, Davidson et al., 2023, Landsverk, Sandquist et al., 2020).

**Figure 5.**
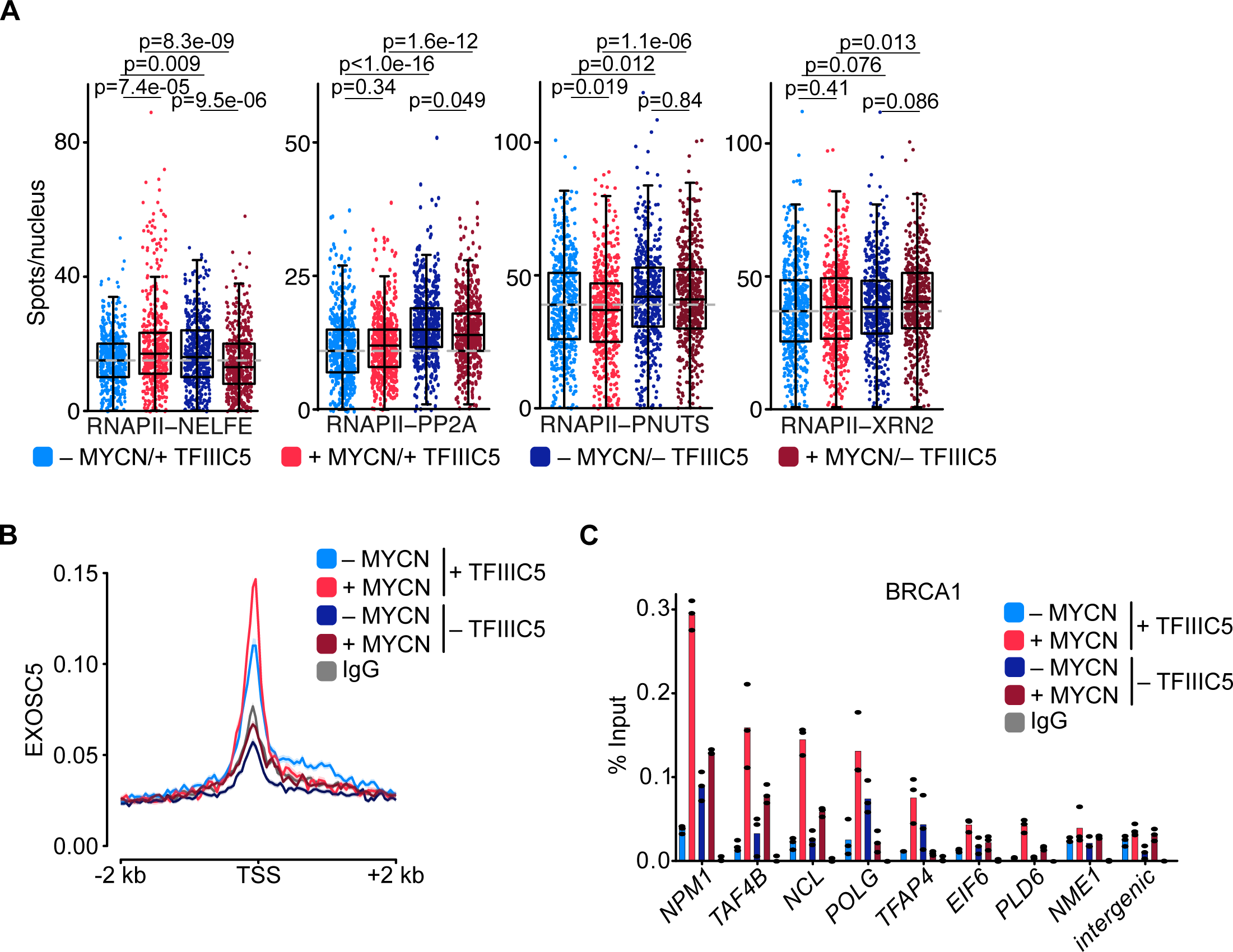
TFIIIC is required for promoter association of the exosome and of BRCA1. **A.** Boxplots showing the number of proximity ligation assay (PLA) signals between RNAPII and NELFE, PP2A, PNUTS or XRN2. SH-EP-MYCN-ER cells were treated with 1 µg/ml Dox (“– TFIIIC5”, 48 h) and/or 4-OHT (“+ MYCN”). EtOH was used as control. For clarity purposes, 500 cells pooled from different replicates were plotted. p-values were calculated comparing the PLA signal of all cells using unpaired Wilcoxon rank sum test. The grey dotted line indicates the median in the control condition (n = 3). **B.** Density plot of CUT&RUN for EXOSC5 binding (N = 14,704 genes) in SH-EP-MYCN-ER cells expressing a Dox-inducible shRNA targeting *TFIIIC5* treated with 4-OHT. Data show mean ± SEM (shade). **C.** BRCA1 ChIP in SH-EP-MYCN-ER cells expressing a Dox-inducible shRNA targeting *TFIIIC5* treated with 4-OHT (4 h). Shown is the mean of technical triplicates of one representative experiment with identical results (n = 2).

Key factors in degradation of aberrant nascent RNAs include XRN2, a 5’-3’ RNA exonuclease, and the exosome, a 3’-5’ exonuclease RNA complex (Cortazar, Erickson et al., 2022, Gerlach, Garland et al., 2022, Noe Gonzalez, Blears et al., 2021). We have previously shown that MYCN can recruit the nuclear exosome to its target promoters (Papadopoulos et al., 2022). MYCN can also recruit BRCA1, which in turn stimulates binding of the mRNA decapping enzyme DCP1 that initiates RNA degradation (Herold et al., 2019). No significant effects were observed in the proximity between RNAPII with the XRN2 exonuclease (Figure 5A). In contrast, CUT&RUN experiments using antibodies directed EXOSC5, a core structural subunit of the exosome (Kilchert, Wittmann et al., 2016), showed that depletion of TFIIIC5 strongly decreased the association of EXOSC5 with regions around the TSS, both in control and in MYCN-expressing cells (Figure 5B). To test whether MYCN-dependent recruitment of BRCA1 depends on TFIIIC, we performed ChIP experiments at multiple MYCN-bound promoters (Figure 5C and Figure S5E). Consistent with our previous observations, only low amount of BRCA1 were found at core promoter in SH-EP cells and that induction of MYCN recruited BRCA1 to all tested promoters (Herold et al., 2019). Depletion of TFIIIC5 had variable effects on the presence of BRCA1 by itself, but abrogated BRCA1 recruitment by MYCN at all promoters tested (Figure 5C and Figure S5E), demonstrating that TFIIIC5 is required for MYCN-dependent recruitment of BRCA1 to multiple target sites. Collectively, the data show that TFIIIC is required for association of the nuclear exosome and of BRCA1 with active promoters and that these effects are enhanced in cells expressing MYCN, arguing that they can account for the enhanced dependence of MYCN expressing cells on TFIIIC. A model summarizing our data is shown in Figure 6.

**Figure 6.**
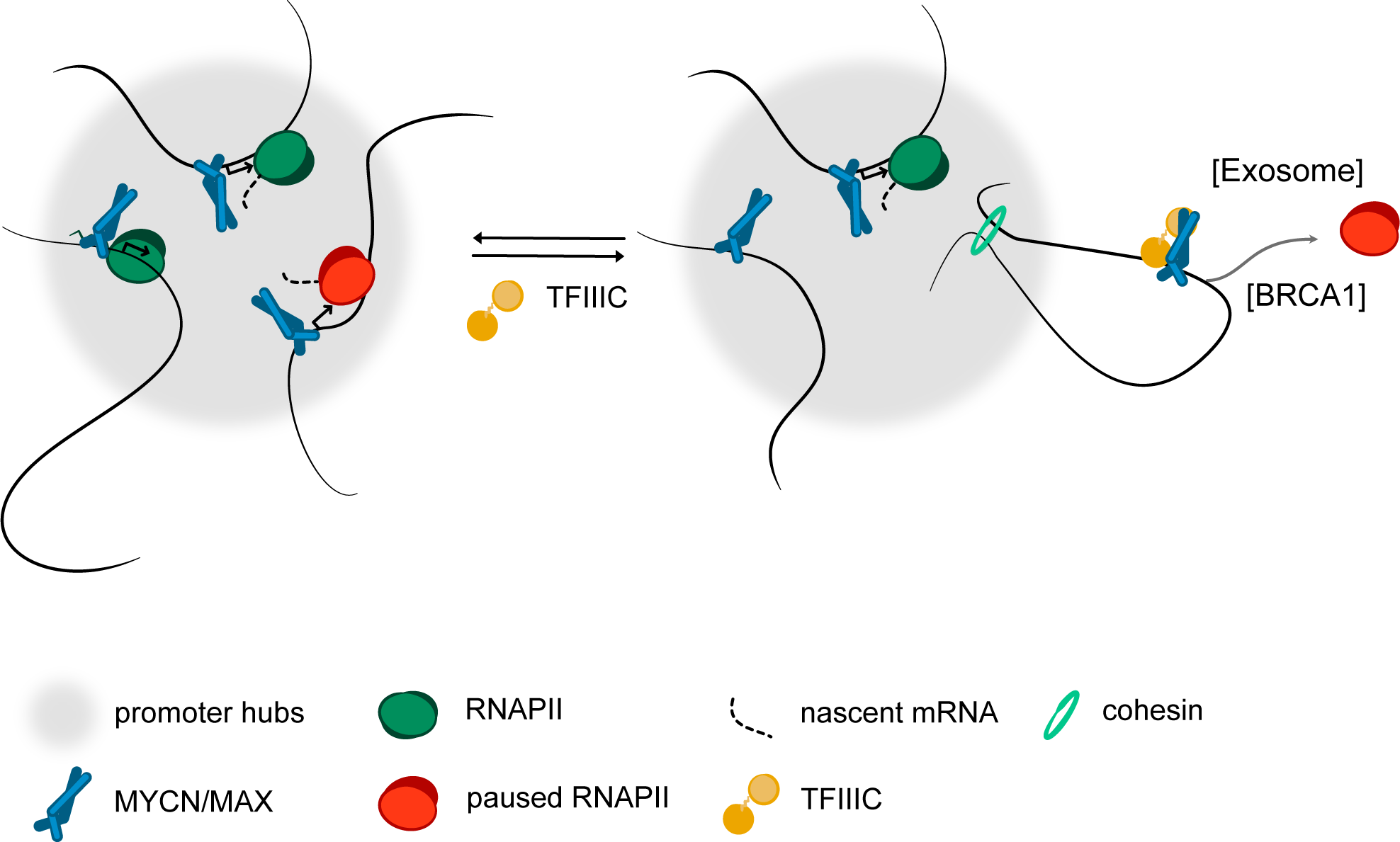
Model. Model summarizing our findings. We propose that complex formation with the TFIIIC complex antagonize the localization of MYCN in promoter hubs and that this enables access of the nuclear exosome and BRCA1 to promoters with paused or stalled RNAPII. Both the exosome and BRCA1 have been implicated in fostering the degradation of nascent RNA at promoters. The precise mechanisms by which MYCN and TFIIIC limit accumulation of non-phosphorylated RNAPII at promoters remain to be determined.

## Discussion

One of the most puzzling aspects of MYC biology is the apparent discrepancy between the global association of MYC and MYCN oncoproteins with open chromatin, which includes virtually all active promoters, particularly in tumor cells, and their much more restricted and often weak effects on the expression of downstream target genes. Most MYC/N binding sites on chromatin do not appear to be functional with respect to gene expression and transcriptional elongation (Kress et al., 2015, Sabo, Kress et al., 2014, Walz et al., 2014). This raises the possibility that such binding sites are indeed non-functional (Pellanda, Dalsass et al., 2021). Alternatively, MYCN has functions in transcription that go beyond gene regulation. In support of the later model, we and others have uncovered critical roles of MYC and MYCN in maintaining the genomic stability of tumor cells, for example, by coordinating transcription with DNA replication and by enabling promoter-proximal double-strand break repair (Baluapuri et al., 2020, Papadopoulos et al., 2023). This raises the question of which direct interaction partners of MYC and MYCN mediate these effects and what the underlying mechanisms are. Here we show that TFIIIC binds directly to the amino-terminus of MYCN, and that a MYCN/TFIIIC complex globally limits the accumulation of non-phosphorylated RNAPII at promoters.

Several mechanisms can account for these observations: First, the amino-terminus of MYC proteins is a transcription-activating domain that can interact with cyclin T1 and CDK8 (Eberhardy & Farnham, 2002) and potentially other co-activators, hence TFIIIC may block MYCN interaction with co-activators that recruit RNAPII to promoters. Since MYCN/TFIIIC also restricts RNAPII accumulation at promoters that are not activated by MYCN, this is unlikely to be the sole mode of action. Second, MYCN and TFIIIC binding sites are often downstream of the transcription start sites and the three-dimensional structures formed by MYCN and TFIIIC may sterically restrict elongation and favour termination of non-phosphorylated RNAPII. Third, TFIIIC is an architectural protein complex and re-localization of promoters may be central to the mechanism of MYCN/TFIIIC action. Active promoters coalesce at a discrete number of sites in the eucaryotic nucleus (Lim & Levine, 2021, Palacio & Taatjes, 2022). Many of the factors involved in basal transcription undergo liquid-liquid phase separation, arguing that transcription initiation takes place in condensate-like structures established by transcription-regulatory domains and by RNAPII itself (Hnisz, Shrinivas et al., 2017, Lewis, Das et al., 2023, Lu, Yu et al., 2018). Consistent with their ubiquitous role in transcription, MYC proteins form condensates by themselves and enter condensates formed by the RNAPII-associated mediator complex (Boija, Klein et al., 2018, Solvie, Baluapuri et al., 2022, Yang, Chung et al., 2022). As described in the introduction, TFIIIC can restrict the localization of genes to sites of active transcription. We show here that MYCN/TFIIIC complexes, in contrast to MYCN, are not part of hubs of active promoters, leading us to propose that TFIIIC antagonizes MYCN participation in promoter hubs and limits the accumulation of paused or stalled RNAPII at promoters (Figure 6). Since TFIIIC competes with the Aurora-A kinase for binding of MYCN and Aurora-A in turn promotes elongation by RNAPII, we propose that the competition between both complexes enables transcriptional elongation on promoters that are activated by MYCN (Buchel et al., 2017, Roeschert, Poon et al., 2021).

As described above, depletion of TFIIIC has no discernible role in elongation and changes in polyA+ mRNA levels, raising the question of what the function of the association might be. The fate of nascent RNA is controlled by several interrelated processes leading to splicing and polyadenylation of the full-length transcript on the one hand, and mis-splicing, premature termination and RNA degradation on the other and MYCN has been implicated in both processes (Noe Gonzalez et al., 2021, Papadopoulos et al., 2022, Schmid & Jensen, 2019). Depletion of TFIIIC strongly decreased the association of the nuclear exosome, an RNA exonuclease complex that degrades multiple forms of aberrant nascent RNA (Schmid & Jensen, 2019), and of BRCA1, which in turn recruits the mRNA decapping enzyme DCP1 at promoters (Herold et al., 2019). The association of both the exosome and BRCA1 with promoters were enhanced in MYCN expressing cells, arguing that they are likely to reflect - at least in part - the activity of the TFIIIC/MYCN complex. At the same time, TFIIIC3 depletion caused splicing errors, which occurred predominantly in downstream exons and were independent of MYCN, arguing that the effect on splicing may reflect an activity of TFIIIC that is independent of its function at promoters. We propose that the TFIIIC/MYCN complex formation exerts a censoring and quality control function for RNAPII at promoters that have not received a full complement of activating signals and hypothesize that this contributes to focus the transcription machinery and metabolic resources on the genes that drive the growth of MYCN-driven tumors.

## Acknowledgements

We thank Vanessa Luzak and Yordan Sbirkov for initial experiments on chromatin architecture. This work was funded by grants from German Cancer Aid (Mildred Scheel Early Career Center, #70113303; G.B.), German Cancer Aid (#70113870; M.E.), German Research Foundation (DFG) (INST 93/1023-1-FUGG and EI222/21-1; M.E.), Alex’s Lemonade Stand Foundation (Crazy 8 Initiative; M.E. and G.B.), Wilhelm Sander-Stiftung (#2023.002.1; S.H.) and Medical Research Council (MR/V029975/1; E.L. and R.B.).

## Author Contributions

R.V., E.L., S.H., M.M., D.F., I.R., D.P. and G.B. performed experiments; C.S.-V. analyzed PLA assays; R.V.. L.U. and P.G. analyzed sequencing data; C.P.A. generated high-throughput data; G.B., R.B., and M.E. devised and supervised experiments and G.B. and M.E. wrote the paper.

## Corresponding Authors

Correspondence should be addressed to Gabriele Büchel and Martin Eilers.

## Data and materials availability

The following published datasets were used: MYCN ChIP-seq and RNA-seq: GSE111905; 4sU-seq: GSE164569. Reviewers can access data from this work under GEO accession GSE223058.

## Methods

### Cell culture

Cell line derived from human neuroblastoma (SH-EP; CVCL_RR78) was verified by STR profiling and grown in RPMI-1640 (Thermo Fisher Scientific). Murine neuroblastoma cells (NHO2A) were grown in RPMI-1640. Murine NIH-3T3 (CVCL_0594) cells were grown in DMEM (Thermo Fisher Scientific). Media were supplemented with 10% fetal calf serum (Capricorn Scientific GmbH) and penicillin/streptomycin (Sigma-Aldrich). All cells were routinely tested for mycoplasma contamination. Inhibitors were used in the following concentrations: MLN8237: 1 µM, 4 h; DRB: 100 µM, 2 h. Growth curves were obtained using the Incucyte® Live-Cell Analysis System.

### Transfection and lentiviral infection

For lentivirus production, HEK293TN (CVCL_UL49) cells were transfected using PEI (Polyethyleneimine, Sigma-Aldrich). Lentiviruses expressing a shRNA targeting *GTFIIIC5* (targeting sequence #1: AAGCGCAGCACCTACAACTACA, #2: TTGATAAATCTTGGCATCTGGG) were produced by transfection of pINDUCER11 (Sequence #1) and pLT3GEPIR (Sequence #2) plasmid together with the packaging plasmid psPAX.2 and the envelope plasmid pMD2.G into HEK293TN cells. Virus-containing supernatant was harvested 24 h and 48 h after transfection. SH-EP-MYCN-ER cells were infected with lentiviral supernatants in the presence of 4 µg/ml polybrene (Sigma-Aldrich) for 24 h. Cells were sorted for GFP and RFP expression. shRNA expression was stimulated by addition of doxycycline (1 µg/ml) for 12 h and cells were FACS-sorted for RFP-GFP double-positive cells. Lentiviruses expressing a shRNA targeting *GTFIIIC2 or GTFIIIC3* (targeting sequence *GTFIIIC2:* TGAAGCAGAAGAATGGTCTGGA*, GTFIIIC3:* TTCATCATTTTCTTGGTTTCAC) were produced by transfection of and pLT3GEPIR plasmid together with the packaging plasmid psPAX.2 and the envelope plasmid pMD2.G into HEK293TN cells. After harvesting supernatant and infection SH-EP-MYCN-ER cells were selected using puromycin and sorted for GFP expression.

For experiments, cells were harvested 48 h after induction with doxycycline (1 µg/ml) or ethanol as control. For induction of the MYCN chimera cells were treated with 4-OHT (200 nM, 4 h) as indicated.

### Constructs

All constructs were cloned from human cDNA, or subcloned from plasmids containing specific human cDNA sequences. All TFIIIC subunit containing complexes (TFIIIC/3xFLAG-MYCN; τA/3xFLAG-MYCN; τA) were expressed using the MultiBac system. For τA/3xFLAG-MYCN, TFIIIC3, TFIIIC5, Strep-TFIIIC6, and 3xFLAG-MYCN aa 2-137, were cloned into pACEBAC1, pIDC, pIDK, and pIDS respectively. These were sequentially assembled into a single vector using Cre-Lox recombination. For the TFIIIC/3xFLAG MYCN complex, 10xHis-TFIIIC1, TFIIIC3, TFIIIC5, TFIIIC6, and 3xFLAG MYCN aa 2-137 were all cloned into pACEBAC1. A single pACEBAC1 vector containing all of these inserts was generated by repetitive sub-cloning of the subunits. The acceptor pACEBAC1 was opened using I-CeuI and SpeI. The region of interest was removed from the donor pACEBAC1 clone using I-CeuI and AvrII. The region of interest was then ligated into the acceptor pACEBAC1. An internal AvrII site had to be silently removed from TFIIIC3 by site directed mutagenesis prior to sub-cloning. The pACEBAC1 with these five members was then used in successive Cre-Lox recombination reactions with pIDK TFIIIC4 and then pIDC TFIIIC2 in order to generate a construct with all seven members of the complex. For the τA complex 6xhis-TEV-TFIIIC3 was cloned into pACEBAC1. This was Cre-loxed with pIDK Strep-TFIIIC6, and pIDC TFIIIC5. MultiBac constructs were validated by restriction enzyme digest and sequencing. Post assembly, constructs containing multiple subunits were validated by analytical PCR and restriction enzyme digest.

### Protein expression and purification

For *Sf*9 expression, bacmid DNA was generated by transformation of expression vectors into *E. coli* DH10MultiBac cells. Purified bacmid DNA (1.6 – 8.3 µg) was transfected into a monolayer of *Sf*9 cells, in a T25 flask, using X-tremeGENE™ HP DNA Transfection Reagent (Roche). After 7 days at 28 °C the P1 virus stock was collected. 1 ml of P1 virus was used to infect 300 ml of *Sf*9 cells at a density of 2 x10^6^ cells/ml. The infected cells were grown for 72 h at 120 rpm and 27 °C prior to harvesting by centrifugation at 1,000 xg for 10 min. The supernatant was used to infect large scale cultures of *Sf*9 cells. 100 ml of virus stock was used per litre of *Sf*9 cells at a density of 1.5 x10^6^ cells/ml. These were harvested after 48 h. *E.coli* expression was performed using BL21(DE3)RIL cells. 10 ml from overnight cultures were used to seed each Litre of LB media (supplemented with 50 μg/ml kanamycin and 35 μg/ml chloramphenical). The cells were grown at 37 °C and 220 rpm until mid log phase (∼0.6 OD_600_) whereupon expression was induced using 0.6 mM final isopropyl β-D-1-thiogalactopyranoside (IPTG). Cells were grown overnight at 20 °C, and were harvested by centrifugation at 6,000 xg for 15 minutes. Cell pellets were stored at -80 °C prior to purification.

For the purification of TFIIIC/3xFLAG-MYCN and τA/3xFLAG-MYCN complexes *Sf*9 pellets were resuspended in a lysis buffer (25 mM Tris pH8, 200 mM NaCl, EDTA free cOmplete PI (Roche), 10 µg/ml DNAseI) then lysed by sonication. Lysate was clarified at 40,000 xg for 20 min. Clarified lysate was applied to 2 ml bed volume Anti-FLAG M2 Affinity Gel (Sigma-Aldrich). The slurry was rotated in the cold room for two hours prior to application to a gravity flow column. Flow was collected and resin washed as follows. For the TFIIIC/3xFLAG-MYCN complex the resin was washed with 40 ml of wash buffer (25 mM Tris pH8, 200 mM NaCl) in four 10 ml batches. For the τA/3xFLAG-MYCN complex the resin was washed with two 10 ml batches of wash buffer, followed by 10 ml of wash buffer with a final NaCl concentration of 350 mM, followed by 10 ml of wash buffer with a final NaCl concentration of 400 mM. The resin was finally washed with 10 ml of wash buffer. In both cases the protein was eluted by addition of 17.5 ml wash buffer spiked with a final concentration of 0.25 mg/ml 3xFLAG peptide. For the τA/3XFLAG-MYCN complex the eluate fractions were spiked with 2 mM final β-mercaptoethanol and then concentrated to less than 1 ml with a Vivaspin 6 10 kDa MWCO centrifugal concentrator. Size Exclusion Chromatography (SEC) was performed using a Superose 6 10/300 GL column attached to an ÄKTA pure FPLC system (Cytiva). The SEC buffer (25 mM Tris pH8, 200 mM NaCl, 3 mM DTT) was flowed over the column at 0.3 ml/min and fractions collected every 0.25 ml over the peaks. In order to do an MYCN alone SEC run, the MYCN peak fractions from several SEC runs of the complex were combined and re-concentrated down to 300 µl. This was then run as previous. For all purified proteins, protein concentration was determined by absorbance at 280 nm. Unless otherwise stated proteins were aliquoted, flash frozen in liquid N_2_ and stored at -80 °C prior to use.

For the purification of τA alone, *Sf*9 pellets were resuspended in a base buffer (25 mM Tris pH 7.4, 2.7 mM KCl, 287 mM NaCl, 2 mM β-mercaptoethanol) supplemented with 0.1 mg/ml DNase I, and 1 EDTA free cOmplete PI (Roche) tablet per 30 ml of buffer. Cells were lysed by sonication, then the lysate clarified at 40,000 xg for 25 min. Clarified lysate was applied to 2.5 ml bed volume His-select Cobalt Affinity Gel (Sigma-Aldrich). The slurry was rotated in the cold room for an hour prior to application to a gravity flow column. The resin was washed sequentially as follows: 50 ml base buffer, 10 ml ATP buffer (25 mM Tris pH 7.4, 137 mM NaCl, 2.7 mM KCl, 5 mM MgCl_2_, 5 mM ATP), 50 ml base buffer with 5 mM imidazole, 50 ml base buffer with 10 mM imidazole. Protein was eluted sequentially by addition of 25 ml of base buffer with 150 mM imidazole followed by 10 ml of the same buffer with 500 mM imidazole. Eluted fractions were pooled and 0.7 mg of his-tagged TEV NIa was added. The mixture was dialysed overnight, using 10 kDa MWCO Snakeskin dialysis tubing, against 4 l of buffer (25 mM Tris pH 7.4, 137 mM NaCl, 2.7 mM KCl, 2 mM β-mercaptoethanol). Post-dialysis the TEV NIa processed complex was rebound to Cobalt resin as per previous. The flow fraction was collected and concentrated. SEC was performed as per the τA/3XFLAG-MYCN complex. τA was concentrated, as per pre-SEC, down to 16.8 μM and used for τA/MYCN complex reconstruction immediately.

Protein purification was assessed by a combination of Coomassie stained SDS-PAGE gels and immunoblotting (TFIIIC90: A301-239A, AB_890667; TFIIIC5: A301-242A, AB_890669; TFIIIC102: A301-238A, AB_890671, Bethyl Laboratories; TFIIIC110: sc-81406, AB_2115237; MYCN: sc-53993, AB_831602, Santa Cruz Biotechnologies; TFIIIC35: NBP2-31851, AB_2891101; TFIIIC1: NBP2-14077, AB_2891102, Novus Biologicals). The SEC elution position of molecular weight standards were established by using Bio-Rad gel filtration standards (cat #1511901). 5 mg/ml BSA was additionally used as the 66 kDa marker. Dextran blue was used to establish the void position of the column.

Aurora-A aa 122-403 D274N C290A C393A was purified as previously described (Bayliss, Sardon et al., 2003). The final buffer was 20 mM Hepes pH 7.5, 150 mM NaCl, 5 mM MgCl_2_, 10% glycerol, 1 mM β-mercaptoethanol. *E. coli* expressed MYCN aa 1-137 pellets were initially disrupted using base buffer (pH 7.5, 100 mM KH_2_PO_4_, 10 mM Tris, 300 mM NaCl, 2 mM β-mercaptoethanol) which was supplemented with 30 mg of lysozyme, 0.3 μg DNAseI, and 0.9 μM MnCl_2_ per 30 ml of base buffer. This was sonicated 15 seconds on 15 seconds off for 150 seconds at 40 % amplitude. The lysate was initially clarified by centrifugation for 30 min at 30,000 xg. As this construct of MYCN expresses into inclusion bodies, the protein had to be recovered from the pellet using urea. All urea solutions were prepared immediately prior to use. The pellet was disrupted by vigorous pipetting after addition of 30 ml of 1 M urea in base buffer followed by sonication as previous. This solution was clarified by centrifugation for 20 min at 30,000 xg. This protocol of disruption of the pellet, following addition of urea containing base buffers, followed by clarification was repeated sequentially for solutions containing 2 M and 4 M urea. 6xHis-tagged MYCN aa 1-137 was found in the soluble fraction of the 4 M urea solution. This was diluted 1/4 with 25 mM Tris pH 7.4, 2.7 mM KCl, 137 mM NaCl, 2 mM β-mercaptoethanol prior to application of the protein to His-Select Cobalt affinity gel (Sigma-Aldrich). The slurry was rotated for 60 min prior to application to a gravity flow column. The resin was washed sequentially as follows with imidazole containing buffer (25 mM Tris pH 7.4, 2.7 mM KCl, 137 mM NaCl, 2 mM β-mercaptoethanol): 50 ml 0 mM imidazole, 50 ml 5 mM imidazole, 25 ml 10 mM imidazole, 25 ml 150 mM imidazole, and 10 ml 500 mM imidazole. Pure fractions were pooled, along with 0.7 mg of his-tagged TEV NIa, and were dialysed overnight using 3.5 kDa MWCO snakeskin dialysis tubing against 4 l of buffer (25 mM Tris pH 7.4, 2.7 mM KCl, 137 mM NaCl, 2 mM β-mercaptoethanol). Post TEV Nia processing, the protein was re-applied to His-Select Cobalt affinity gel (Sigma-Aldrich). The flow through fractions were pooled and concentrated (Vivaspin Turbo 3,000 MWCO) prior to SEC. SEC was performed using an ÄKTA prime and 16/600 S75 column. Pure fractions were pooled and concentrated as per pre SEC. Protein concentration was determined using absorbance at 280 nm.

### τA/3xFLAG-MYCN native mass spectrometry

The τA/3xFLAG-MYCN complex was desalted into 200 mM ammonium acetate using 75 μl Zeba spin desalting columns (ThermoFisher Scientific). Samples were analysed by nanoelectrospray ionisation MS using a quadrupole-orbitrap MS (Q-Exactive UHMR, ThermoFisher Scientific) using gold/palladium coated nanospray tips prepared in-house. The MS was operated in positive ion mode using a capillary voltage of 1.3 kV, capillary temperature of 250 °C and S-lens RF of 200 V. In source trapping was used with a desolvation voltage of - 100 V for 4 µs. Extended trapping was not used. The quadrupole mass range was 2000-15000 m/z. Nitrogen gas was used in the HCD cell with a trap gas pressure setting of 5. Orbitrap resolution was 12500, detector m/z optimisation was low. 5 microscans were averaged. Mass calibration was performed by a separate injection of sodium iodide at a concentration of 2 µg/µl. Data processing was performed using QualBrowser 4.2.28.14 and deconvoluted using UniDec (Marty, Baldwin et al., 2015).

### τA/3xFLAG-MYCN intact mass spectrometry

Protein desalting and mass analysis was performed by LC-MS using an M-class ACQUITY UPLC (Waters UK, Manchester, UK) interfaced to a Xevo QToF G2-XS mass spectrometer (Waters UK, Manchester, UK). Samples were diluted to 1 µM using 0.1% TFA. 1 µl of the 1 µM sample was run on an Acquity UPLC Protein BEH C4 column (300 Å, 1.7 µm, 2.1 mm × 100 mm, Waters UK) with an Acquity UPLC Protein BEH VanGuard Pre-Column (300 Å, 1.7 µm, 2.1 mm × 5 mm, Waters UK). Buffer A was 0.1% formic acid in water, and buffer B was 0.1% formic acid in AcN (v/v basis). System flow rate was kept constant at 50 µl/min. Protein sample was loaded on to the trap column in 20% acetonitrile/0.1% formic acid and washed for 5 min. Following valve switching, the bound protein was eluted by a gradient of 20-95% solvent B in A over 10 min. The column was subsequently washed with 95 % solvent B in A for 5 min before re-equilibration at 20% solvent B in A ready for the next injection. The mass spectrometer was calibrated using a separate injection of glu-fibrinopeptide. Data were processed using MassLynx 4.2.

### Protein identification/Peptide LC-MS analysis

Gel bands were excised and chopped into small pieces (∼ 1 mm^3^), covered with 30% ethanol in a 1.5 ml microcentrifuge tube and heated to 56 °C for 30 min with shaking. The supernatant was removed, replaced with fresh ethanol solution and was again heated to 56 °C for 30 min with shaking. This was repeated until all Coomassie stain was removed from the gel. The gel slices were then dehydrated by covering with 100% acetonitrile and left for five minutes before the supernatant was discarded and replaced with a fresh aliquot of acetonitrile. Proteins were reduced by adding 100 µl 20 mM DTT solution before incubation at 57 °C for 1 h with shaking. The supernatant was removed and once the gel pieces were at room temperature proteins were alkylated by adding 100 µl 55 mM iodoacetic acid. The samples were then incubated at room temperature in the dark for 30 min with shaking. After removing the supernatant, the gel slices were then covered with 100% acetonitrile and left for five minutes. The acetonitrile was removed, and the gel pieces were left to dry in a laminar flow hood for 60 min. Once dry, the gel slices were cooled on ice then they were covered with ice-cold protease solution and left on ice for 20 min to rehydrate. Trypsin solution (20 ng/µl in 25 mM ammonium bicarbonate) was added to the unknown bands, chymotrypsin solution (25 ng/µl in 25 mM ammonium bicarbonate). Excess protease solution was removed, and the gel slices were covered with a minimal amount of 25 mM ammonium bicarbonate. After briefly vortexing and centrifuging, the gel slices were incubated at 37 °C while shaking for 18 h. The resulting digest was vortexed and centrifuged. The supernatant was recovered and added to a 1.5 ml tube containing 5 µl acetonitrile/ water/ formic acid (60/35/5; v/v). 50 µl acetonitrile/ water/ formic acid (60/35/5; v/v) was added to the gel slices and vortexed for an additional 10 min. The supernatant was pooled with the previous wash and one additional wash of the gel slices was performed. The pool of three washes was dried by vacuum centrifugation. The peptides were reconstituted in 20 µl 0.1% aqueous trifluoroacetic acid prior to analysis. 4 µl sample were injected onto an in house-packed 20 cm capillary column (inner diameter 75 µm, 3.5 µm Kromasil C18 media). An EasyLC nano liquid chromatography system was used to apply a gradient of 4–40% ACN in 0.1% formic acid over 45 min at a flow rate of 250 nl/min. Total acquisition time was 60 min including column wash and re-equilibration. Separated peptides were eluted directly from the column and sprayed into an Orbitrap Velos Mass Spectrometer (ThermoFisher Scientific, Hemel Hempstead, UK) using an electrospray capillary voltage of 2.7 kV. Precursor ion scans were acquired in the Orbitrap with resolution of 60,000. Up to 20 ions per precursor scan were selected for fragmentation in the ion-trap. Dynamic exclusion of 30 s was used. Peptide MS/MS data were processed with PEAKS Studio X+ (Bioinformatic Solutions Inc, Waterloo, Ontario, Canada) and searched against the Uniprot databases for *H. sapiens* and *S. frugiperda* proteins (release 2021_02). Carbamiodomethylation was selected as a fixed modification, variable modifications were set for oxidation of methionine and deamidation of glutamine and asparagine. MS mass tolerance was 15 ppm, and fragment ion mass tolerance was 0.3 Da. The peptide false discovery rate was set to 1%.

### Immunoblots

Whole cell extracts were prepared using RIPA buffer (50 mM HEPES pH 7.9, 140 mM NaCl, 1 mM EDTA; 1% Triton X-100, 0.1% sodium deoxycholate, 0.1% SDS) containing protease and phosphatase inhibitor cocktails (Sigma-Aldrich). Lysates were cleared by centrifugation and protein concentrations were determined by Bradford.

Protein samples were separated on Bis-Tris gels and transferred to a PVDF membrane (Millipore). For immunoblots showing multiple proteins with similar molecular weight, one representative loading control is shown. Vinculin (VCL) or GAPDH were used as loading control. Antibodies used in this study: MYCN (B8.4.B): sc-53993, AB_831602, Santa-Cruz Biotechnology; TFIIIC5: A301-242A, AB_890669, Bethyl Laboratories; MYC (Y69): ab32072, AB_731658, Abcam; VCL (h-VIN1): V9131, AB_477629, Sigma-Aldrich; GAPDH: 2118, AB_561053, Cell Signaling; TFIIIC2: sc-81406, AB_2115237, Santa-Cruz; TFIIIC3: sc-393235, Santa-Cruz.

### Proximity Ligation Assay (PLA)

SH-EP-MYCN-ER cells were plated in 384-well plates (PerkinElmer), treated with Dox and/or 4-OHT and fixed with methanol for 20 min. After blocking for 30 min with 5% BSA in PBS, cells were incubated over night at 4 °C with primary antibodies: Total RNA Polymerase II (F12): sc-55492, (N20): sc-899, Santa-Cruz Biotechnology; TFIIIC5: A301-242A, Bethyl Laboratories; NELFE: ABE48, Merck; PP2A: 2038, Cell Signaling; PNUTS: A300-439-1, Bethyl Laboratories; XRN2: A301-103A, Bethyl Laboratories. PLA was performed using Duolink® In Situ Kit (Sigma-Aldrich) according to the manufactureŕs protocol. Nuclei were counterstained using Hoechst 33342 (Sigma-Aldrich). Pictures were taken with Z-stacks of four planes at 0.5 µm distance at 40-fold magnification in Operetta® CLS High-Content Imaging System. Analysis was performed in Harmony® High Content Imaging and Analysis Software. Thirty image fields per well were acquired with a total of at least 500 cells per sample.

### High-throughput sequencing

ChIP and ChIP-seq was performed as described previously (Roeschert et al., 2021). For each ChIP or ChIP-Rx seq experiment, 5x10^7^ cells per immunoprecipitation condition were fixed for 5 min at room temperature with formaldehyde (final concentration, 1%). Fixation was stopped by adding 125 mM glycine for 5 min. Cells were harvested in ice-cold PBS containing protease and phosphatase inhibitors (Sigma-Aldrich). As exogenous control (spike-in), murine NHO2A or NIH3T3 cells were added at a 1:10 cell ratio during cell lysis. Cell lysis was carried out for 20 min in lysis buffer I (5 mM PIPES pH 8.0, 85 mM KCl, 0.5% NP-40) and nuclei were collected by centrifugation (1,500 rpm, 10 min, 4 °C). Crosslinked chromatin was prepared in lysis buffer II (10 mM Tris pH 7.5, 150 mM NaCl, 1 mM EDTA, 1% NP-40, 1% sodium deoxycholate, 0.1% SDS) and fragmented by using the Covaris Focused Ultrasonicator M220 for 50 min per ml lysate. For each IP reaction, 15 µl (for ChIP) or 100 μl (for ChIP-seq) Dynabeads Protein A and Protein G (Thermo Fisher Scientific) were pre-incubated overnight with rotation in the presence of 5 mg/ml BSA and 3 µg (for ChIP) or 10-15 μg (for ChIP-seq) antibody (MYCN (B8.4.B): sc-53993, AB_831602; TFIIIC5: A301-242A, AB_890669; RNAPII pSer2: ab5095, AB_304749; RNAPII (8WG16): sc-56767, AB_785522; BRCA1: A300-000A, AB_67367, Bethyl Laboratories). Chromatin was added to the beads, and IP was performed for a minimum of 6 h at 4 °C with rotation. Beads were washed three times each with washing buffer I (20 mM Tris pH 8.1, 150 mM NaCl, 2 mM EDTA, 1% Triton X-100, 0.1% SDS), washing buffer II (20 mM Tris pH 8.1, 500 mM NaCl, 2 mM EDTA, 1% Triton X-100, 0.1% SDS), washing buffer III (10 mM Tris pH 8.1, 250 mM LiCl, 1 mM EDTA, 1% NP-40, 1% sodium deoxycholate; including a 5 min incubation with rotation) and TE buffer. Chromatin was eluted twice by incubating with 200 µl elution buffer (100 mM NaHCO_3_, 1% SDS in TE) for 15 min with rotation. Input samples and eluted samples were de-crosslinked overnight. Protein and RNA were digested with proteinase K and RNase A, respectively. DNA was isolated by phenol-chloroform extraction followed by an ethanol precipitation and analyzed by qPCR using StepOnePlus Real-Time PCR System (Thermo Fisher Scientific) and SYBR Green Master Mix (Thermo Fisher Scientific) or sequencing on the Illumina Next-Seq 2000.

After DNA extraction occupancy of different proteins were assessed by RT-PCR. Primers were used for *TFAP4*(forward: CCGGGCGCTGTTTACTA; reverse: CAGGACACGGAGAACTACAG), *POLG* (forward: CTTCTCAAGGAGCAGGTGGA; reverse: TCATAACCTCCCTTCGACCG), *NPM1* (forward: TTCACCGGGAAGCATGG; reverse: CACGCGAGGTAAGTCTACG), Intergenic region (forward: TTTTCTCACATTGCCCCTGT; reverse: TCAATGCTGTACCAGGCAAA), *NCL* (forward: CTACCACCCTCATCTGAATCC; reverse: TTGTCTCGCTGGGAAAGG), *NME1* (forward: GGGGTGGAGAGAAGAAAGCA; reverse: TGGGAGTAGGCAGTCATTCT), *PLD6* (forward: GCTGTGGGTCCCGGATTA; reverse: CCTCCAGAGTCAGAGCCA), *TAF4B* (forward: AAGGTCGTCGCTCACAC, reverse: GCGTGGCTATATAAACATGGCT), *RPL22* (forward: CCGTAGCTTCCTCTCTGCTC, reverse: ACCTCTTGGGCTTCCTGTCT), *CCND2* (forward: GCCAGCTGCTGTTCTCCTTA, reverse: CCCCTCCTCCTTTCAATCTC). Shown analysis of RT-PCR show mean of technical triplicates as well as an overlay of each data point to indicate the distribution of the data.

For ChIP- or ChIP-Rx-seq, DNA was quantified using the Quant-iT PicoGreen dsDNA assay (Thermo Fisher Scientific). DNA library preparation was done using the NEBnext Ultra II DNA Library Prep Kit (New England Biolabs) following manufacturer’s instructions. Quality of the library was assessed on the Fragment Analyzer (Agilent) using the NGS Fragment High Sensitivity Analysis Kit (1-6,000 bp; Agilent). Finally, libraries were subjected to cluster generation and base calling for 75 cycles on Illumina NextSeq 500 platform.

CUT&RUN followed by sequencing was performed as described in Meers et al. (Meers, Tenenbaum et al., 2019). 5 x10^5^ cells were harvested using Accutase (Sigma-Aldrich). Cells were washed in wash buffer (20 mM HEPES pH 7.5, 150 mM NaCl, 0.5 mM Spermidine). After washing, cells were coupled to ConA-coated magnetic beads (Polysciences Europe), permeabilized and subsequently incubated with the primary antibody (EXOSC5: NBP2-14952, 1:100, Novus Biologicals) in antibody binding buffer (wash buffer, 0.05% digitonin, 2 mM EDTA) over night at 4 °C. Cells were washed in Dig-Wash buffer (Wash buffer, 0.05% digitonin) and MNase (700 ng/ml) was added to each sample for 1 h at 4 °C. After washing with Dig-wash buffer and Low-Salt Rinse buffer (20 mM HEPES pH 7.5, 0.5 mM Spermidine, 0.05% digitonin) incubation buffer (3.5 mM HEPES pH 7.5, 10 mM CaCl_2_, 0.05% digitonin) was added for 30 min at 4 °C. STOP buffer (170 mM NaCl, 20 mM EGTA, 0.05% digitonin, 20 µg/ml RNase A, 25 µg/ml Glycogen) was added to stop the MNase digestion. DNA fragments were released at 37 °C for 30 min. Decrosslinking was performed for 1 h at 50 °C after adding 0.01% SDS and 10 mg/ml Proteinase K followed by Phenol Chloroform extraction. Precipitated DNA was resuspended in 30 µl 0.1x TE buffer. For library preparation NEBnext Ultra II DNA Library Prep Kit (New England Biolabs) was used. Pre-PCR samples were purified using AmpureXP beads with a ratio of 1.75x. Then the eluted material was used with 16 PCR cycles. The libraries were cleaned using Agentcourt AMPure XP Beads (ratio 0.8x, Beckman Coulter), quality, quantity and fragment size assessed on the Fragment Analyzer (Agilent) using the NGS Fragment High Sensitivity Analysis Kit (1-6,000 bp; Agilent). Finally, libraries were subjected to cluster generation and base calling for 75 cycles on Illumina NextSeq 500 platform.

Phosphorylated linker Hi-C (pLHi-C) was based on Hi-C (Dixon, Selvaraj et al., 2012) and *in situ* Hi-C (Rao, Huntley et al., 2014). The main differences consisted in replacing the digestion enzyme and the Klenow fill-in step to increase resolution and overall protocol efficiency. 1x10^6^ cells per condition were fixed with formaldehyde (1% final concentration) for 10 min at room temperature (RT). Reaction was stopped by adding 125 mM glycine for 15 min at RT. Cells were harvested with ice-cold PBS supplemented with protease and phosphatase inhibitors (Sigma-Aldrich). Nuclei isolation was performed by adding buffer I (10 mM Tris-HCl pH 8, 10 mM NaCl, 0.5% NP-40, 1x protease inhibitors; Sigma-Aldrich) coupled with douncer homogenization. The resulting solution was centrifuged (2,500 xg, 5 min, 4 °C), the nuclei pellet was washed with 1x NEBuffer 3.1 (NEB) and incubated in buffer IIa (0.2% SDS, 1x NEBuffer 3.1; NEB) for 10 min at 50 °C. Buffer IIb (2% Triton X-100, 1.5 x NEBuffer 3.1, 1x protease inhibitors;) was added to a final volume of 943 µl. Sample was incubated for 15 min at 37 °C. 850 U DpnII (NEB) and 35 µl rSAP (NEB) were added and sample was incubated for at least 8 h at 37 °C. Modified DNA oligos, 5’-GATCCCCAAATCT-3’ and 5’-GATCAGAT[BtndT]TGGG-3’ with 5’ end phosphate (Sigma-Aldrich), were annealed (81 µl of each Oligo 100 µM, 1x T4 ligase buffer; NEB) at 98 °C for 5 min, followed by 20 min at RT. After digestion, sample was washed with 1x T4 ligase buffer (NEB) (2,500 xg, 5 min, 4 °C). Nuclei pellet was resuspended in buffer IIIa (0.3% SDS, 1x T4 ligase buffer) and incubated for 15 min at 60 °C. Buffer IIIb (3% Triton X-100, 1x T4 ligase buffer, 1x protease inhibitors; NEB, Sigma-Aldrich) was added to a final volume of 1 ml and sample was incubated for 15 min at 37 °C. The nuclei pellet was resuspended in buffer IV (160 µl annealed oligos, 1.2x T4 ligase buffer, 6,000 U T4 ligase, 1x protease inhibitors; NEB) to a final volume of 800 µl after washing it with 1x T4 ligase buffer (NEB) (2,500 xg, 5 min, 4 °C). Sample was incubated overnight at 16 °C. rSAP in the digestion followed by phosphorylated linker in ligation was introduced to decrease the likelihood of self-ligation, increasing the final yield. Nuclei pellet was isolated by centrifugation (2,500 xg, 5 min, 4 °C) and mixed with buffer V (10 mM Tris-HCl pH 7.6, 1 mM EDTA, 0.1% SDS,1x protease inhibitors). Chromatin fragmentation was performed using the Covaris Focused Ultrasonicator M220 for 30 min per ml lysate. Samples were de-crosslinked overnight. Protein and RNA were digested with proteinase K and RNase A, respectively. DNA was isolated by ChIP DNA Clean & Concentrator (Zymo Research) according to the manufacturer’s instructions. Biotin pull-down was performed with MyOne Streptavidin C1 beads (Thermo Fisher Scientific) according to manufacturer’s instructions. Library preparation was performed on beads using NEBNext ChIP-Sequencing Library Prep Master Mix Set for Illumina (NEB). The manufacturer’s instructions were followed, apart from introducing a washing step between each module of the protocol with buffer VI (3x, 5 mM Tris-HCl pH 8, 0.5 mM EDTA, 1 M NaCl, 0.05% Tween) followed by one with 1x TE buffer. Quality of the library was assessed on the Fragment Analyzer (Agilent) using the NGS Fragment High Sensitivity Analysis Kit (1-6,000 bp; Agilent). Libraries were subjected to cluster generation and base calling for 150 cycles paired end on Illumina NextSeq 500 platform.

Spike-in phosphorylated linker HiChIP (spLHiChIP) and phosphorylated linker HiChIP (pLHiChIP) were based on pLHi-C, ChIP-Rx, and the original HiChIP protocol (Mumbach et al., 2016). spLHiChIP and pLHiChIP benefit from the same modules of pLHi-C fixation, digestion, ligation, chromatin fragmentation, and library preparation. The difference consists in the ChIP module, which follows the exact same steps from IP of target protein to de-crosslinked overnight stated in the ChIP workflow. The ChIP module allocated between the chromatin fragmentation and library preparation in pLHi-C workflow defines the spLHiChIP and pLHiChIP workflows. spLHiChIP differs from pLHiChIP by adding murine NHO2A cells as an exogenous control (spike-in) at a 1:20 cell ratio during nuclei isolation. This allows normalization by murine paired end valid tags in a similar fashion as murine reads for ChIP-Rx. 10x10^6^ cells per condition were fixed with formaldehyde (1% final concentration) for 10 min at room temperature (RT). Reaction was stopped by adding 125 mM glycine for 15 min at RT. Cells were harvested with ice-cold PBS supplemented with protease and phosphatase inhibitors (Sigma-Aldrich). Nuclei isolation was performed by adding buffer I (10 mM Tris-HCl pH 8, 10 mM NaCl, 0.5% NP-40, 1x protease inhibitors; Sigma-Aldrich) coupled with douncer homogenization. The resulting solution was centrifuged (2,500 xg, 5 min, 4 °C), the nuclei pellet was washed with 1x NEBuffer 3.1 (NEB) and incubated in buffer IIa (0.2% SDS, 1x NEBuffer 3.1; NEB) for 10 min at 50 °C. Buffer IIb (2% Triton X-100, 1.5 x NEBuffer 3.1, 1x protease inhibitors;) was added to a final volume of 943 µl. Sample was incubated for 15 min at 37 °C. 850 U DpnII (NEB) and 35 µl rSAP (NEB) were added and sample was incubated for at least 8 h at 37 °C. Modified DNA oligos, 5’-GATCCCCAAATCT-3’ and 5’-GATCAGAT[BtndT]TGGG-3’ with 5’ end phosphate (Sigma-Aldrich), were annealed (81 µl of each Oligo 100 µM, 1x T4 ligase buffer; NEB) at 98 °C for 5 min, followed by 20 min at RT. After digestion, sample was washed with 1x T4 ligase buffer (NEB) (2,500 xg, 5 min, 4 °C). Nuclei pellet was resuspended in buffer IIIa (0.3% SDS, 1x T4 ligase buffer) and incubated for 15 min at 60 °C. Buffer IIIb (3% Triton X-100, 1x T4 ligase buffer, 1x protease inhibitors; NEB, Sigma-Aldrich) was added to a final volume of 1 ml and sample was incubated for 15 min at 37 °C. The nuclei pellet was resuspended in buffer IV (160 µl annealed oligos, 1.2x T4 ligase buffer, 6,000 U T4 ligase, 1x protease inhibitors; NEB) to a final volume of 800 µl after washing it with 1x T4 ligase buffer (NEB) (2,500 xg, 5 min, 4 °C). Sample was incubated overnight at 16 °C. Nuclei pellet was isolated by centrifugation (2,500 xg, 5 min, 4 °C) and mixed with buffer V (10 mM Tris-HCl pH 7.6, 1 mM EDTA, 0.1% SDS,1x protease inhibitors). Chromatin fragmentation was performed using the Covaris Focused Ultrasonicator M220 for 30 min per ml lysate. For each IP reaction, 100 μl Dynabeads Protein A and Protein G (Thermo Fisher Scientific) were pre-incubated overnight with rotation in the presence of 5 mg/ml BSA and 10 μg antibody (MYCN (B8.4.B): sc-53993, AB_831602; TFIIIC5: A301-242A, AB_890669). Chromatin was added to the beads, and IP was performed for a minimum of 6 h at 4 °C with rotation. Beads were washed three times each with washing buffer I (20 mM Tris pH 8.1, 150 mM NaCl, 2 mM EDTA, 1% Triton X-100, 0.1% SDS), washing buffer II (20 mM Tris pH 8.1, 500 mM NaCl, 2 mM EDTA, 1% Triton X-100, 0.1% SDS), washing buffer III (10 mM Tris pH 8.1, 250 mM LiCl, 1 mM EDTA, 1% NP-40, 1% sodium deoxycholate; including a 5 min incubation with rotation) and TE buffer. Chromatin was eluted twice by incubating with 200 µl elution buffer (100 mM NaHCO_3_, 1% SDS in TE) for 15 min with rotation. Input samples and eluted samples were de-crosslinked overnight. Protein and RNA were digested with proteinase K and RNase A, respectively. DNA was isolated by phenol-chloroform extraction followed by an ethanol precipitation. DNA library preparation was done using the NEBnext Ultra II DNA Library Prep Kit (New England Biolabs) following the manufacturer’s instructions. Quality of the library was assessed on the Fragment Analyzer (Agilent) using the NGS Fragment High Sensitivity Analysis Kit (1-6,000 bp; Agilent). Finally, libraries were subjected to cluster generation and base calling on Illumina NextSeq 500 platform.

RNA sequencing was performed by extracting RNA with RNeasy Mini Kit (Qiagen) according to manufacturer’s instructions. On-column DNase I digestion was performed followed by mRNA isolation with the NEBNext Poly (A) mRNA Magnetic Isolation Kit (NEB). Library preparation was done with the Ultra II Directional RNA Library Prep for Illumina following the manufacturer’s manual. Libraries were size selected using SPRIselect Beads (Beckman Coulter) after amplification with 9 PCR cycles. Library quantification and size determination was performed with the Fragment Analyzer (Agilent) using the NGS Fragment High Sensitivity Analysis Kit (1-6,000 bp; Agilent). Libraries were subjected to cluster generation and base calling for 100 cycles paired end on Illumina NextSeq 2000 platform.

### Bioinformatics analysis and statistics

All libraries were subjected to NextSeq 500 or NextSeq 2000 (Illumina) sequencing following the manufacturer’s guidelines. Base call data was converted and demultiplex to FASTQ files by bcl2fastq Conversion Software v1.1.0 (Illumina). Quality control of each dataset was done using FastQC tool.

ChIP-seq and CUT&RUN sequencing reads were aligned to human (hg19/ GRCh37 assembly) using Bowtie2 v2.3.5.1. with default parameters (Langmead & Salzberg, 2012). Normalization by aligned read number was performed on the input and on all correspondent IP samples. For each sample, the number of spike-in normalized reads was calculated by dividing the number of reads mapping exclusively to hg19 by the number of reads mapping exclusively to mm10 and multiplying this number with the smallest number of reads mapping to mm10 of all samples. After normalization, files were converted to bedGraph with “bedtools genomecov” v2.26 (Quinlan & Hall, 2010). These files were used to plot genome browser track examples via the package plotgardener v1.012 in R v4.1.1.

MACS2 v2.1.2 (Zhang, Liu et al., 2008) was used for peak calling with the following ChIPseq-datasets & parameters, with inputs serving as control: MYCN (GSM3044606, GSM3044608; dup = 5, q = 1x10^-3^), TFIIIC5 in SH-EP-MYCN-ER (this work; dup = 5, q = 10^-3^), and RAD21 (this work; dup = 5, p = 10^-2^). H3K4me1, H3K4me3, and H3K27ac peaks were called with SICER v1.1 (Xu, Grullon et al., 2014), using input as control and the parameters: redundancy threshold = 1, window size = 200 bp, fragment size = 75 bp, effective genome fraction = 0.74, gap size = 600, FDR = 0.001. Peak calling results were refined and confirmed by visual inspection.

RNA sequencing reads were aligned to human (hg19/ GRCh37 assembly) using STAR aligner v2.7.9a with the default parameters and gene quantification (--quantMode GeneCounts) (Dobin, Davis et al., 2013). The output table with the gene counts was loaded into R v4.1.1. and processed with DESeq2 package accordingly to their manual (Love, Huber et al., 2014). Genes with or more than 15 counts in at least 3 samples or more were selected, and all samples were normalized by size factors. The fold average for 3 independent biological replicates was calculated on treatment to control as showed in the correspondent figure and legend. The result was averaged in 100 or 150 bins for a total of 14,085 genes and displayed as dot plot. Alternative splicing events were identified using rMATS (Shen et al., 2014b).

pLHi-C, pLHiChIP and spLHiChIP sequencing reads were processed in two steps. In the first step they were all processed in the same way, HiC-Pro v 2.11.4 (Servant, Varoquaux et al., 2015) trims the reads for the linker sequence, aligns it to hg19 and filters it by DpnII restriction fragments of the same genome assembly. The default parameters were used for the alignment steps, except for the length of the seed (-L) parameter that was set to 25 for the global options and 15 for the local one, and mismatch (-N) that was set to 1. Read trimming was performed by inserting combinations of the linker sequence (GATCAGATTTGGGGATC, GATCCCCAAATCTGATC) in the ligation site parameter. Reads considered as duplicates were accepted and multi-mapped reads were only removed for pLHiChIP and spLHiChIP. Quality control for all steps until all valid pairs (<u>p</u>aired <u>e</u>nd tags; PETs) output was also performed thru HiC-Pro. We compared HiC-Pro’s standard quality controls example (IMR90 replicate 1 sample; GSM862724), according to Servant *et al*. 2015 (Servant et al., 2015), to the base of our methods variation, pLHi-C.

In the second step, spLHiChIP samples undergo the same workflow described for the first step replacing hg19 for mm10 assembly. PETs from mm10 were removed from hg19 PETs and a spike-in normalization was applied to hg19 PETs in the same way explained for ChIP-Rx hg19 aligned reads. DNA loop calling was performed on spLHiChIP and pLHiChIP PETs using hichipper v0.7.7 (Lareau & Aryee, 2018). ChIP-seq peak calling sites were used as pre-determined peaks. All the default parameters with a loop maximum distance (--max-dist) of 5,000,000 were used for all proteins. We compared the DNA loops calling quality control for Oct4 HiChIP (GSM2238510) from the original method description (Mumbach et al., 2016) to our method variation for MYCN and TFIIIC5. An extra quality control to show enrichment and specificity was performed by plotting MYCN pLHiChIP, its input pLHi-C and its ChIP-seq with pLHiChIP and pLHi-C normalized to the same number of PETs. DNA loops calling results were refined and confirmed by visual inspection. The mango output files from the loop calling were then loaded in R, filter by a q-value equal or smaller than 0.01 and used in all further plots and analyses. All examples showing DNA loops, ChIP-seq and annotation tracks in different genomic regions/locus were plotted using plotgardener package in R environment.

The different genome functional annotations were retrieved from UCSC table browser (Karolchik, Hinrichs et al., 2004). Promoters, gene bodies and transcription end sites (TES) come from Reference Sequence (RefSeq) collection of genes for hg19 assembly and were adapted via R language. For RNAPII targets, promoters were defined as TSS ± 0.5 kb (Buchel et al., 2017), gene bodies as the sum of exons and introns, TES as TES ± 0.3 kb. Enhancers were defined as in Walz *et al*. (2014) (Walz et al., 2014): H3K3me1 and H3K27ac overlapping sites without H3K4me3. Overlapping and discontinuous regions were obtained by using peak calling outputs as inputs in intersectBed (BEDTools) with standard parameters.

Overlapping between different pLHiChIP was calculated by “bedtools pairtopair” function (BEDTools) using the options to prevent self-hits and ignore strands. The type of overlap was set to report when both anchors of the different pLHiChIP overlaps (-type either). This provided outputs of loops with and without overlapping that could be used for other analyses. For discrimination between the different types of overlaps, the type both was removed from the type either and the two types were then reported. Overlaps were called joint sites and adapted as Venn diagrams by centering the coordinates and numbers to MYCN anchors or MYCN-TFIIIC5 anchors. TADs overlap corresponds to common sites in at least eight out of nine cell lines reported in (GSE35156) (Rao et al., 2014).

The GenomicInteractions v1.28 (Harmston, Ing-Simmons et al., 2015) package in R language was used to calculate and report the overlaps between the different pLHiChIP anchors and their functional annotations. Overlaps on different functional annotations were reported in the following order of priority: enhancer, promoter, TES, gene body, tRNA, rRNA, miRNA, snRNA, snoRNA, SINE, LINE, LTR, and satellite. The bar graphs reporting the functional annotation of each anchor are proportional to the total number of loops of each pLHiChIP, numbers were rounded and plotted by GenomicInteractions and ggplot2 v3.3.5 packages in R environment. Motif analysis was performed by using the FIMO algorithm (Grant, Bailey et al., 2011) of the MEME Suite software toolkit v5.3.3 (Bailey, Johnson et al., 2015). The function getfasta (BEDTools) extracted the hg19 FASTA sequence from each pLHiChIP anchor to be used independently as input for motif search. The background model was generated by the function fasta-get-markov (MEME Suite) using hg19 FASTA with standard parameters. Pre-defined motifs for MYCN (E-box) (MA0104.4: JASPAR 2018 (Sandelin, Alkema et al., 2004)), B-box, and A-box were scanned using the options “--thresh 1.0 --text” with the default parameters (Chan, Lin et al., 2021, Pavesi, Conterio et al., 1994). Resulting hits were then filtered by p-value <= 1x10^-4^. In R environment, duplicated hits of the same motif in the same anchor were filtered by the one with the lowest p-value. Different motifs in the same anchor window were reported according to the decreasing priority: E-box, B-box, and A-box. The overlap between anchors and motifs was computed by GenomicInteractions package and the corresponding heatmap was plotted in R.

To visualize the influence of the MYCN interaction on gene expression (Fig 1C), MYCN bound genes (as defined by MACS2 on MYCN ChIP-seq data, above) were subdivided into genes that are bound by MYCN without loops (“no MYCN loops”) and genes bound by MYCN forming loops (above, “MYCN loops”). For both classes, RNA-sequencing reads between TSS & TSS+2 kb were counted using “bedtools intersect” on spike-normalized ChIP-seq data for RNAPII (A10), MYCN (GSM3044606), TFIIIC5; processed transcript data were obtained from Büchel *et al*. 2017 (RNA-sequencing, GSE111905 (Buchel et al., 2017)) and Papadopoulos et al. 2021 (4sU-sequencing, GSE164569 (Papadopoulos et al., 2022)), respectively. Plots were generated with “boxplot” and statistical significance calculated using Students’ t-test (unpaired, two-sided, unequal variance), both in R v3.6.3.

Over-representation analysis for Molecular Signatures Database (MsigDB (Subramanian, Tamayo et al., 2005)) were calculated by clusterProfiler package (Wu, Hu et al., 2021) in R environment. Different functional classes of loops were retrieved from the overlaps and discontinuous loops by the GenomicInteractions package. Gene names were converted to HGNC symbol name using AnnotationDbi package and sorted by the number of PETs. Duplicated gene names were merged and the median of their number of PETs was used. A hypergeometric test was calculated on the sorted list for MSigDB collections H (hallmark gene sets) or C5 (ontology gene sets) using a 0.05 q-value threshold.

MYCN spLHiChIP treated sample was normalize by its control counterpart in each biological replicate before plotted as graph bar in R enviroment. The ratio of TFIIIC5 binding (ChIP-Rx) per number of interaction (spLHiChIP) was calculated in R environment v4.1.1. The number of ChIP-Rx reads was calculated to TSS ± 2 kb for all genes using bedtools intersect (BEDtools) with default parameters. The number of spLHiChIP interactions was calculated for the same region. The ratio was calculated on reads to number of interactions per each sample and plotted as bar graph.

Networks were reconstituted by using the R package igraph v1.2.11 and Cytoscape v3.9 (Brockmann, Poon et al., 2013). The different classes of loops were retrieved via the GenomicInteractions package and converted to the simple interaction file format and to a node table. These files were used in Cytoscape for visualization, using the Edge-weighted Spring-Embedded Layout based on the number of PETs for each loop. The resulting MYCN network reports the nodes as functional annotations; explained before. In parallel, the sif file was used as input for the igraph package. This package was used to define the membership of each anchor to the different clusters and to quantify the number of nodes per cluster. The statistical significance comparing the number of nodes per cluster between two groups were calculated via Wilcoxon rank sum test.

Density plots for the different regions were performed using ngs.plot v2.41.3 (Shen, Shao et al., 2014a). Default parameters were used for samples normalized by reads number. For spike-in normalized samples, we used a custom variation of ngs.plot that allows the spike-in normalization by skipping the internal normalization step. All densities plots and metaplots were based on gene expression for SH-EP-MYCN-ER cells ± MYCN induction (4-OHT treatment) – evaluated by RNA-sequencing (GSE111905; Herold *et al*., 2019). Based on this, all expressed genes gene set consists of 14,704 genes, down- and up-regulated gene sets are represented by 613 and 921 genes, respectively. MYCN-down and -up regulated genes are defined by negative and positive fold change upon ±4-OHT treatment of genes with q-values less than 0.05 and expression filter by counts per million. In R v4.1.1., Wilcoxon rank sum test (unpaired, two-sided) was used to compare the different conditions on the plotted region per ChIP.

PLA’s p-values were calculated by comparing the signal of the pool of the cells in all replicates using Wilcoxon rank sum test (unpaired, two-sided) in R v4.1.1.

### Materials, data and code availability

All plasmids generated are available upon request.

## Supplementary Figures

**Figure S1:**
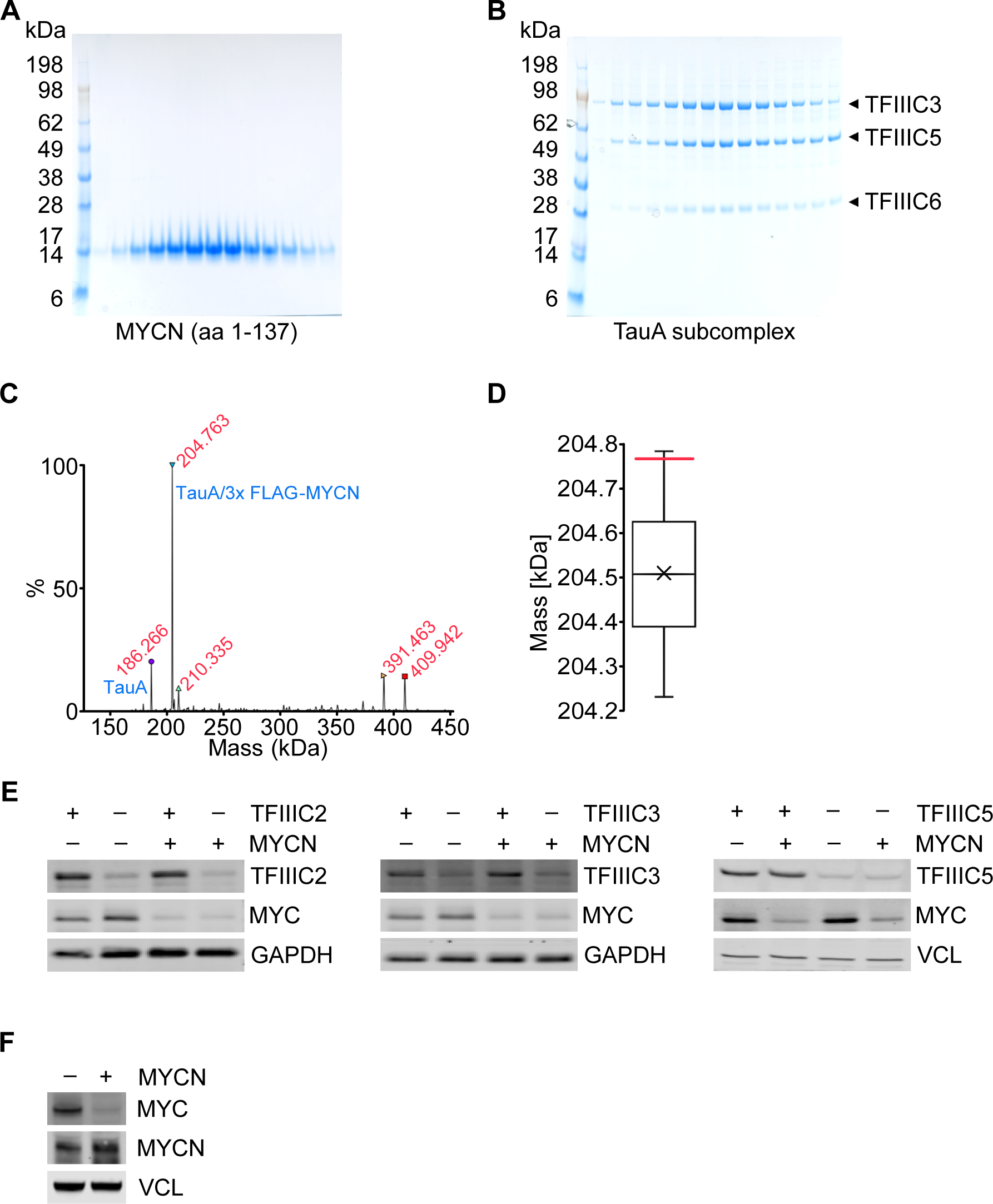
Characterization of MYCN/TFIIIC complexes. **A.** SDS-PAGE gel of recombinant purified MYCN (aa 1-137) expressed in *E. coli* cells. **B.** SDS-PAGE gel of recombinant purified τA subcomplex which was expressed in *Spodoptera frugiperda* (Sf9) cells. **C.** Deconvoluted spectra of native mass spectrometry for the τA/3x FLAG-MYCN complex. Masses are shown in red, identities of complexes are marked in blue. **D.** Box plot of the 30 possible τA/3x FLAG-MYCN complex molecular weights based on the masses observed using intact mass spectrometry. The red line indicates the mass observed for the complex by native mass spectrometry (204,763 Da). **E.** Immunoblot showing levels of TFIIIC2, TFIIIC3 or TFIIIC5 and MYC in SH-EP-MYCN-ER cells expressing doxycycline (Dox)-inducible shRNAs targeting the different *TFIIIC* subunits (n = 3). Where indicated cells were treated with Dox (1 µg/ml, 48 h) and/or 4-OHT (200 nM, 4 h), respectively. EtOH was used as control. In the following panels addition of 4-OHT is indicated by “+ MYCN” and Dox treatment by “– TFIIIC”. **F.** Immunoblot of MYC and MYCN in SH-EP-MYCN-ER cells after induction of MYCN with 4-OHT (200 nM, 4 h). VCL was used as a loading control.

**Figure S2.**
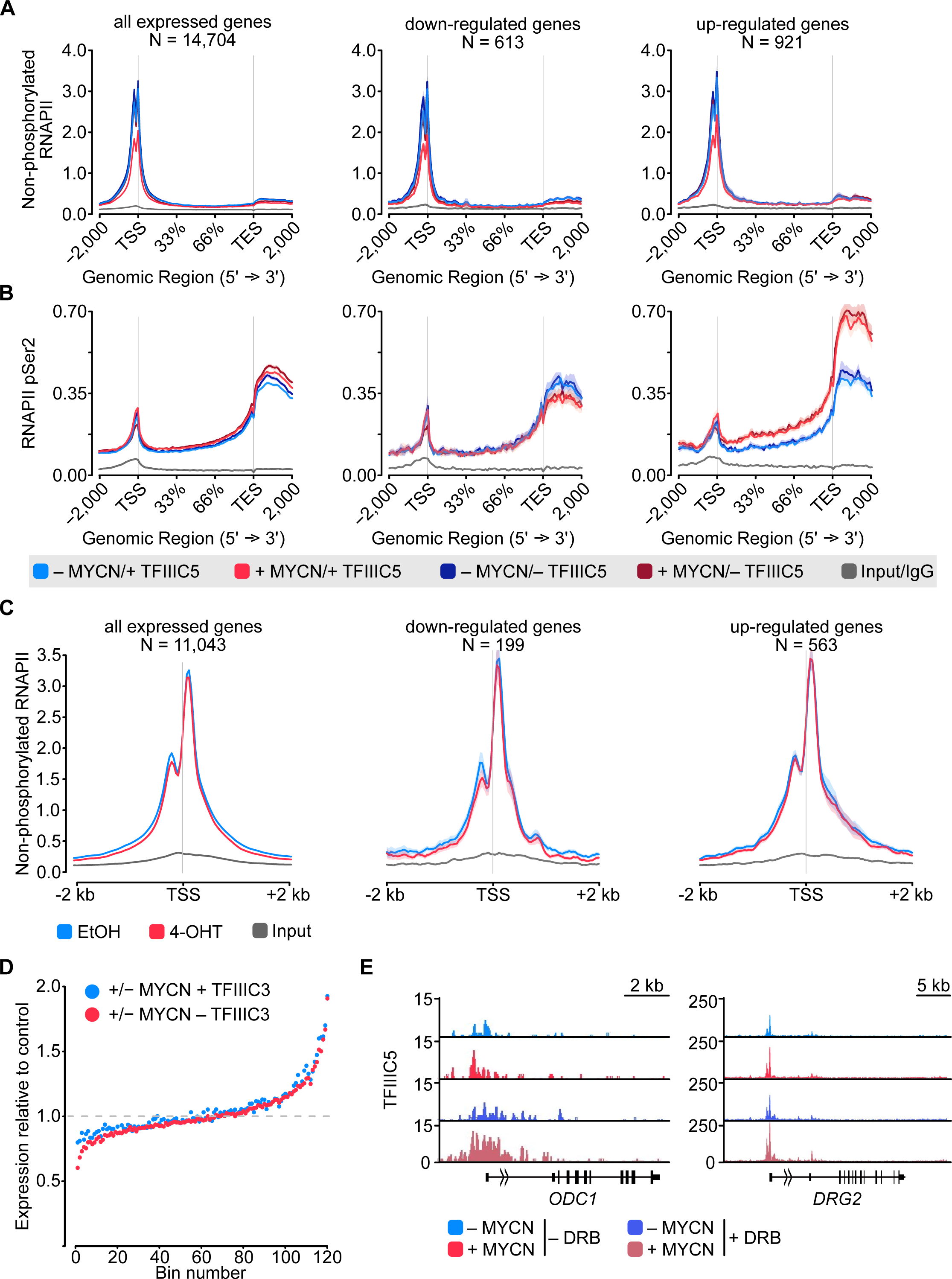
Effects of MYCN and TFIIIC on RNA polymerase II. **A.** Metagene plot of ChIP-Rx signal for non-phosphorylated RNAPII. Data show mean (line) ± standard error of the mean (SEM indicated by the shade) of different gene sets based on an RNA-seq of SH-EP-MYCN-ER cells ± 4-OHT. **B.**. Metagene plot of ChIP-Rx signal for RNAPII pSer2. Data are presented as described in A. **C.** Average density plot of ChIP-Rx signal for non-phosphorylated RNAPII in SH-EP cells treated with 4-OHT. Data show mean (line) ± SEM (indicated by the shade) of different gene sets based on an RNA-seq of SH-EP-MYCN-ER cells ± 4-OHT. The signal is centered to the TSS ± 2 kb (n = 2). **D.** Average bin dot plot for RNA-seq of SH-EP-MYCN-ER showing mRNA expression normalized by control per bin. Cells were co-treated with 1 µg/ml Dox (“– TFIIIC3”, 48 h) and/or 4-OHT (“+ MYCN”, 4 h) as EtOH as control. Expression was normalized by its control. Each bin represents 100 genes of a total of 12,091 genes. Dotted line marks the relative expression at 1 (n = 3). **E.** Browser tracks for TFIIIC5 ChIP-Rx at the indicated gene loci. SH-EP-MYCN-ER cells were treated with DRB and/or 4-OHT, respectively. EtOH was used as control.

**Figure S3:**
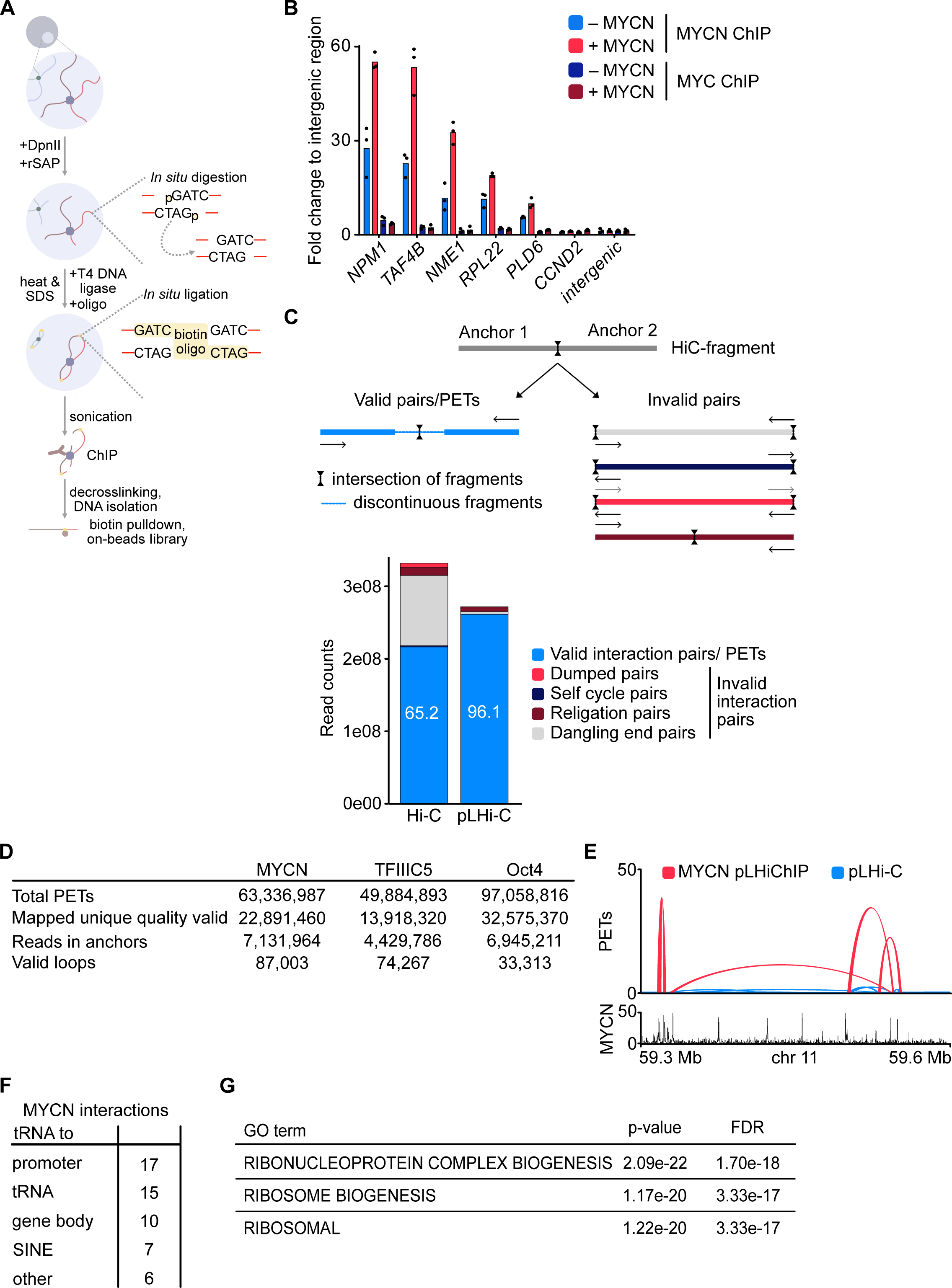
Characterization of HiChip Methods. **A.** Diagram showing the workflow of pLHiChIP. The workflow corresponds to the original HiChIP protocol. For a detailed description, see Methods. **B.** MYCN and MYC ChIP in SH-EP-MYCN-ER cells before and after activation of MYCN with 4-OHT (200 nM, 24 h). Shown is the mean and points of technical triplicates normalized to an intergenic region (n = 2). **C.** (Top) Diagram illustrating the difference between valid and invalid pairs after mapping of spLHiChIP data. Valid pairs harbor non-contiguous elements of chromosomal DNA, whereas invalid pairs harbor contiguous stretches of chromosomal DNA. (Bottom) Bar graph showing percentage of valid and invalid reads in our dataset (right) compared to a published HiC protocol (left). **D.** Table showing quality controls for MYCN and TFIIIC5 spLHiChIP compared to the original Oct4 HiChIP. **E.** Representative example of pLHiChIP track for MYCN (red) and a pLHiC track (blue) showing the strongly increased PETs number of the MYCN interactions after MYCN-immunoprecipitation relative to input (pLHi-C). Data are superimposed with a browser view of a MYCN ChIP-seq. **F.** Table showing interactions involving *tRNA* genes in MYCN pLHiChIP. **G.** Table showing the top three terms, their p-values and the FDR from enrichment analysis for MSigDB C5 collection of all MYCN network.

**Figure S4:**
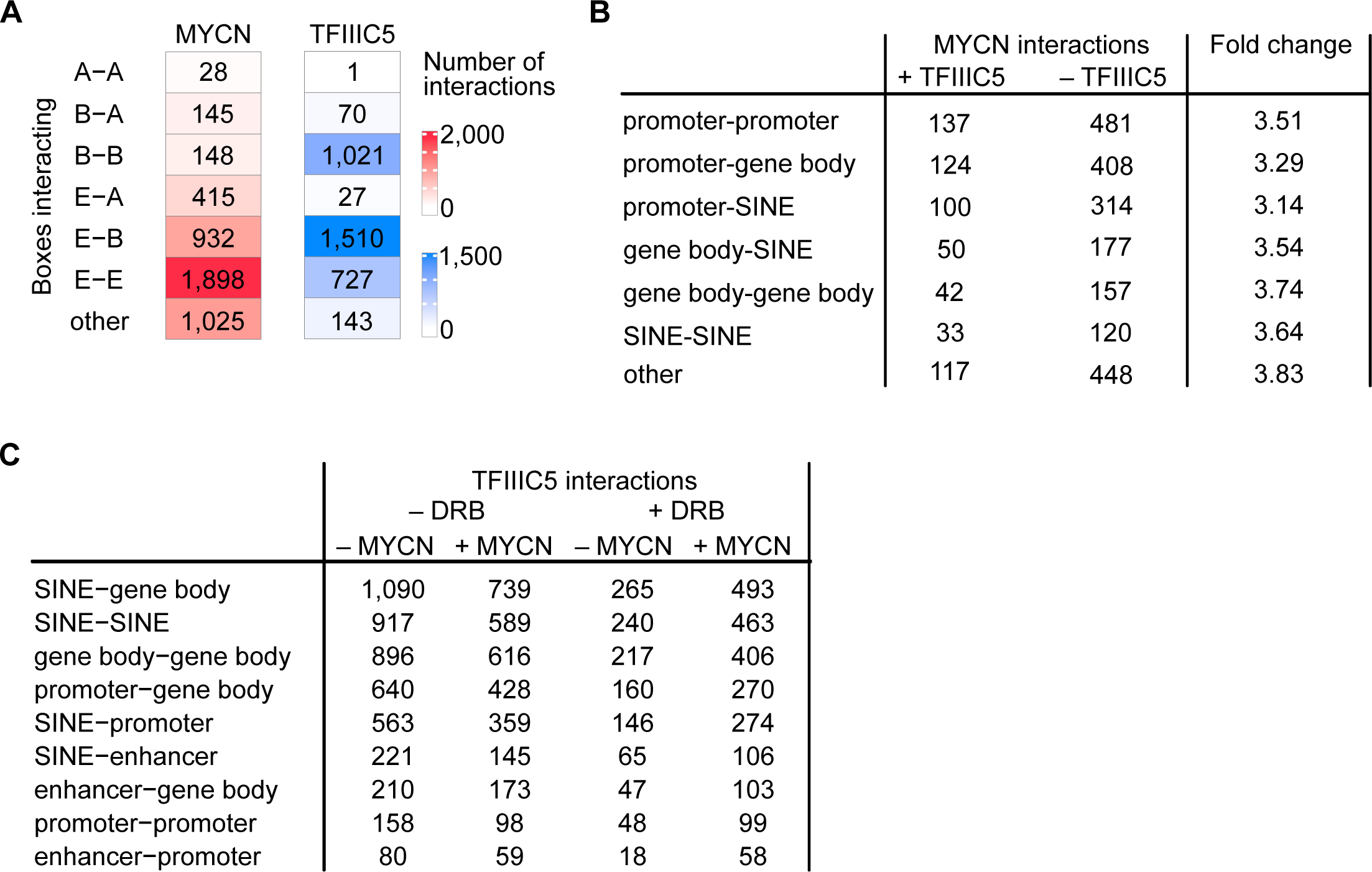
Three-dimensional interactions of MYCN and TFIIIC. **A.** Heatmap showing analysis of sequence motifs characteristic for interacting boxes on both anchors of the interactions. Color reflects the number of interactions. **B.** Table summarizing numbers of MYCN interactions in the presence and absence of TFIIIC5 in SH-EP-MYCN-ER cells and the TFIIIC5-dependent change. Data merged of 2 independent biological replicates. **C.** Table displaying total numbers of TFIIIC5 interactions in SH-EP-MYCN-ER ± MYCN + DRB.

**Figure S5:**
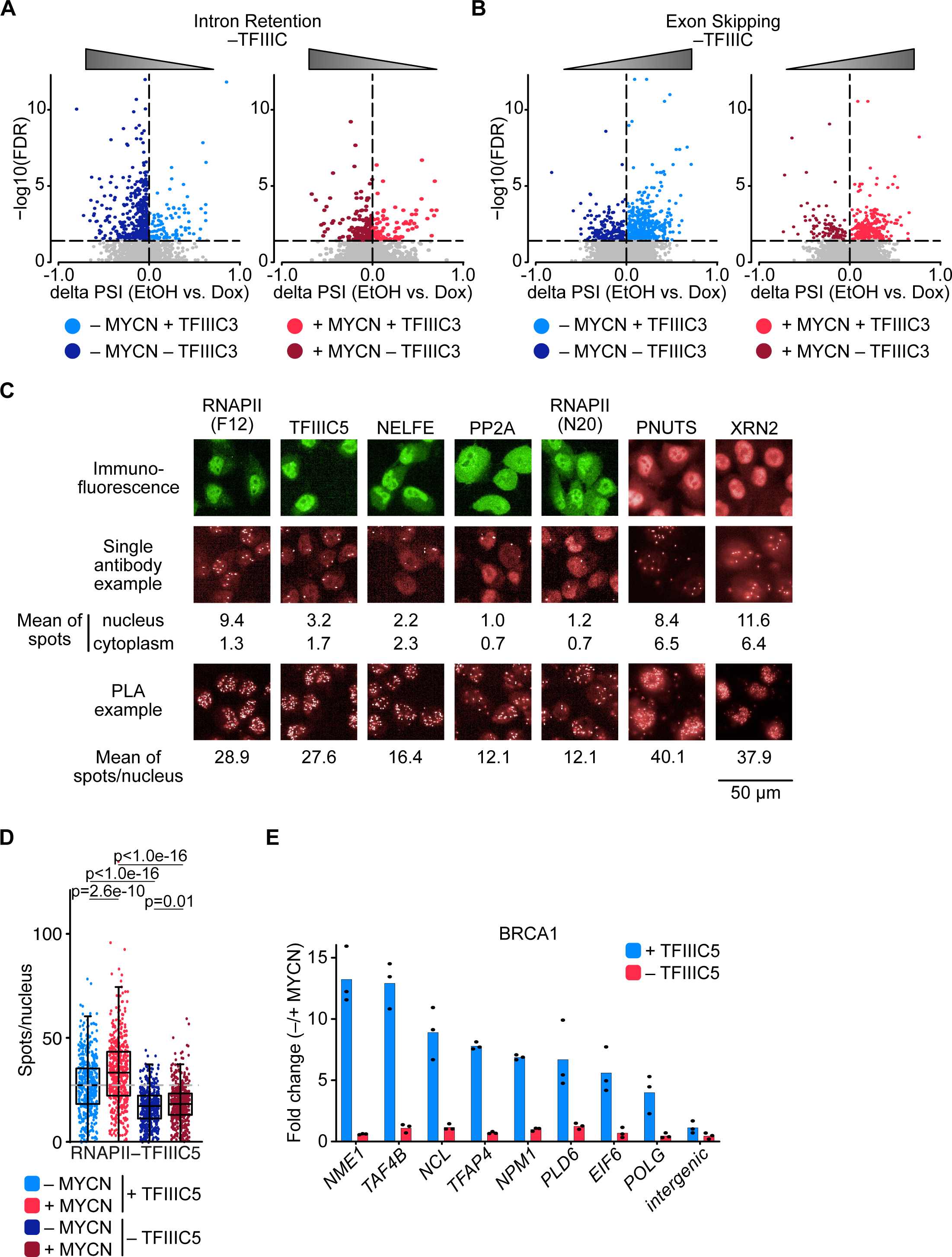
Effects of TFIIIC on splicing and termination factors. **A.** Volcano Plot depicting changes in intron retention upon TFIIIC3 knock-down in the absence (left) and presence (right) of MYCN. Triangle reflects the effect of the splicing error after TFIIIC knock-down. **B.** Volcano Plot depicting changes in exon skipping upon TFIIIC3 knock-down in the absence (left) and presence (right) of MYCN. Triangle reflects the effect of the splicing error after TFIIIC knock-down. **C.** Controls for proximity ligation assays (PLAs) shown in Figure 5A. Immunofluorescence show specificity of the antibody. Single antibodies PLAs were performed in parallel to each PLA. Mean of signal in the nucleus and cytoplasm was calculated and compared to mean of the signal per nucleus of the PLA. **D.** Boxplots showing the number of proximity ligation assay (PLA) signals between RNAPII and TFIIIC5. SH-EP-MYCN-ER cells were treated with 1 µg/ml Dox (“– TFIIIC5”, 48 h) and/or 4-OHT (“+ MYCN”). EtOH was used as control. For clarity purposes, 500 cells pooled from different replicates were plotted. p-values were calculated comparing the PLA signal of all cells using unpaired Wilcoxon rank sum test. The grey dotted line indicates the median in the control condition (n = 3). **E.** BRCA1 ChIP in SH-EP-MYCN-ER cells expressing a Dox-inducible shRNA targeting *TFIIIC5* treated with 4-OHT. Data show fold change of BRCA1 binding after induction of MYCN in the presence (blue) or absence (red) of TFIIIC5 (n = 2).

